# Pre-Border Gene Foxb1 Regulates the Differentiation Timing and Autonomic Neuronal Potential of Human Neural Crest Cells

**DOI:** 10.1101/646026

**Authors:** Alan W. Leung, Francesc López-Giráldez, Cayla Broton, Kaixuan Lin, Maneeshi S. Prasad, Jacqueline C. Hernández, Andrew Z. Xiao, Martín I. Garcia-Castro

## Abstract

What are the factors that are induced during the transitory phases from pluripotent stem cells to lineage specified cells, how are they regulated, and what are their functional contributions are fundamental questions for basic developmental biology and clinical research. Here, we uncover a set of pre-border (pB) gene candidates, including forkhead box B1 (FOXB1), induced during human neural crest (NC) cell development. We characterize their associated enhancers that are bound by pluripotency factors and rapidly activated by β-catenin-mediated signaling during differentiation. Surprisingly, the endogenous transient expression of FOXB1 directly regulates multiple early NC and neural progenitor loci including *PAX7*, *MSX2*, *SOX1*, and *ASCL1*, controls the timing of NC fate acquisition, and differentially activates autonomic neurogenic versus mesenchymal fates in mature NC cells. Our findings provide further insight into the concept of the less characterized pB state and clearly establishes FOXB1 as a key regulator in early cell fate decisions during human pluripotent stem cell differentiation.

## INTRODUCTION

A fundamental question in developmental biology is how pluripotent cells transit from pluripotency towards lineage specification (Kalkan and Smith, 2014). Recent advent of time-series single cell RNA-sequencing has revealed dynamic transcriptional features and cell states during pluripotent cell differentiation *in vivo* and *in vitro* (Pijuan-Sala et al., 2018). Gain of function by overexpression, and loss or reduction of function (knockdown and knockout) approaches, such as CRISPR/CRISPR interference/siRNA/miRNA screening and reporter assays, have also been used to identify genes that are important for initial cell fate decision from pluripotent cells (Arduini and Brivanlou, 2012; Betschinger et al., 2013; Genga et al., 2019; Hackett et al., 2018; Ma et al., 2015; Wang et al., 2012). To functionally characterize novel cell states, there is an urgency to develop experimental pipelines that will integrate the various molecular inputs, effectors, and feedback regulations to infer an increasingly sophisticated road map of cellular differentiation.

Neural crest (NC) is an embryonic cell population that emerges from the dorsal neural tube, migrates extensively, and generates peripheral nervous system neurons and glia, and ectomesenchyme-derived craniofacial bone and cartilage, amongst many other derivatives (Prasad et al., 2019; Stuhlmiller and Garcia-Castro, 2012). This wide range of differentiation potential defies canons of sequential segregation of potential because NC has been considered to emerge from the ectoderm, and bone and cartilage derivatives are normally only associated with mesoderm, not with ectoderm. This issue remains unresolved, with a general perception that the ectodermally-restricted NC lineage acquires ectomesenchymal capacity. Hall suggested that NC constituted a fourth germ layer, but provided little mechanistic evidence (Hall, 2000). Work in chick and rabbit embryos has challenged the classic perception of NC ontogeny, postulating that the early, and thus anterior NC, are specified during gastrulation, independently of mesoderm and or neural interactions and thus postulating a pre-germ layer origin (Basch et al., 2006; Betters et al., 2018). Our unpublished work has further identified in chick blastula embryos with matching experiments in human embryonic stem (ES) cells a NC specification status in pre-streak epiblast within chick embryos and within hours after initiation of NC induction from human ES cells that is distinct molecularly and functionally from pluripotent stem cells (Prasad et al 2019, submitted).

Previously we established a robust model in human pluripotent stem cells, which produce NC cells with anterior character, and allowed us to interrogate the ontogeny of human anterior NC (Leung et al., 2016). In this process of NC induction, we uncovered a class of genes, which we dubbed as pre-border (pB) genes (including *GBX2*, *SP5*, *ZIC3* and *ZEB2*) that are induced prior to, and with distinct signaling requirements than, classic neural plate border genes (Leung et al., 2016). The biological significance of these pB genes is currently not clear. Evidence from vertebrate embryos and cell reprogramming supports the existence of a ‘pB-like’ cell state prior to the emergence of neural border or definitive neurectoderm (Basch et al., 2006; Betters et al., 2018; Thier et al., 2019; Trevers et al., 2018). The concept of pB cell state however is poorly characterized. In our previous study, we proposed the idea that NC specification takes place prior and independently of neurectoderm commitment (Leung et al., 2016). Here we aim to scrutinize the transcriptome and dissect the molecular functions of pB candidate genes and aim to provide further insights into the earliest lineage specification events during the exit of pluripotency towards NC lineage.

Forkhead box (FOX) family proteins are important factors in disease and development (Golson and Kaestner, 2016). They contain a DNA binding domain known as a forkhead or wing helix domain and these proteins can act either as pioneer factors to open local chromatin structures, as classic transcription factors, or both (Iwafuchi-Doi and Zaret, 2016; Lalmansingh et al., 2012). They play essential roles in the regulation of mammalian pluripotency (Krishnakumar et al., 2016; Respuela et al., 2016) and in the specification and further differentiation of neural crest (Kos et al., 2001; Lukoseviciute et al., 2018; Sasai et al., 2001; Seo et al., 2017; Seo and Kume, 2006; Teng et al., 2008; Xu et al., 2018).

Foxb1 is strongly expressed in epiblast, neural plate, neuroepithelium, and midbrain neural folds including NC progenitors (Labosky et al., 1997). Lineage tracing in mice suggests contributions to multiple lineages, including NC derivatives (Zhao et al., 2007). *Foxb1^-/-^* mice displayed developmental delay, posterior truncation, and open neural tube phenotype (Labosky et al., 1997). These mice also lacked a subgroup of medial mammillary body neurons implicated in spatial memory and behaviors (Wehr et al., 1997). Foxb1 could act as a survival factor for these mammillary neurons during the formation of mammillo-thalamic axonal tract (Alvarez-Bolado et al., 2000). In frog embryos, forced expression of Foxb1 led to increased neural induction mediated via cooperation with a Pou transcription factor (Takebayashi-Suzuki et al., 2011) It is of note that a human patient with intellectual disability and distinctive facial features was found with large 15q22.2 deletion spanning the *FOXB1* locus (Yamamoto et al., 2014). The function of FOXB1 during human ES differentiation is currently unknown.

In this work, by studying the transcriptomic changes, enhancer utility, molecular functions, and genomic binding patterns of pB candidates, we aim to reveal the importance of pB intermediate during the exit of pluripotency and the specification of NC cells. Our data uncovers 68 candidates for pB intermediate and reveals that these gene candidates are regulated by elements pre-occupied by pluripotency factors and β-catenin co-factors. Our data also reveals major functions of FOXB1, a key pB candidate, in controlling the timing of NC differentiation, and regulating the differentiation potential of NC precursors towards autonomic neuronal fates. Finally, we provide genomic-binding data of FOXB1 showing that these functions are carried out by direct transcriptional controls on key neurogenic and NC gene loci. Altogether, our data provides important new insights into the poorly characterized pB cellular state and reveals FOXB1 as an essential regulator of human ES cell differentiation. Our data further highlights the critical function of FOXB1 in conferring autonomic neurogenic potential to pre-NC cells.

## RESULTS

### ES cell transcriptional response to canonical WNT signaling during NC induction

Human NC cell induction using our established protocol takes 5 days (Leung et al., 2016) and progresses via a pB and a neural border stage (Fig. 1A) to generate SOX10^+^ NC cells that do not express the definitive neurectoderm marker PAX6 along their course of induction (Fig. 1B) (Leung et al., 2016). To trace the transcriptomic trajectory of early human NC progenitors, we performed time-course mRNA sequencing (RNA-seq) of human ES cells, cells derived from differentiation day 3 (pre-NC, representing a mixture of pB and neural border progenitors) and differentiation day 5 (representing NC progenitors). RNA-seq samples were classified by unsupervised hierarchical clustering according to their differentiation status (Fig. 1C). Differential expression (p<0.05) was respectively detected at 3338 (ES cells versus day 3), 4424 (ES cells versus day 5) and 2009 (day 3 versus day 5) loci. The RPKM values and fold changes comparing different time points for all mapped transcripts are provided in table S1. Factors known to regulate pluripotency and self-renewal including *POU5F1* and *NANOG*, except *SOX2,* and components for FGF (*FGF2*, *DUSP6*), and NODAL (*NODAL*, *GDF3*, *LEFTY1*, *LEFTY2*, *TDGF1*) signaling were downregulated during differentiation (Fig. 1D). Previously identified pB transcripts *GBX2*, *ZIC3*, *ZEB2*, and *SP5*, as well as *MEIS2* (a known NC factor), were dramatically upregulated at day 3 (Fig. 1E). This was followed by a gradual induction of neural border and NC transcripts including *PAX3*, *PAX7*, *TFAP2A*, *NR2F1*, *MSX1*, *MSX2*, *SNAI2, NR2F1*, *CDH6*, *SOX9* and *SOX10*, while *FOXD3* and *ETS1*, displayed a biphasic expression pattern (Fig. 1D, E). As expected for their essential roles in human NC cell induction (Leung et al., 2016), ligands for WNT and BMP pathways, as well as their receptors (*FZDs*) and extracellular regulators (*CHRD*, *NOG*, *BMPER*), were upregulated synchronously during differentiation (Fig. S1A). Signaling components for Hedgehog (*HHIP, GLI1, GLI3, SMO, SUFU*), NOTCH (*JAG1*, *HES3/4*), and insulin growth factor (IGF)(*IGF1*, *IGF1R, IGFBP2/5* and *PAPPA*) pathways were either induced or repressed with their general trends indicative of either a blockade (Hedgehog) or activation (Notch and IGF) of the pathways (Fig. S1B). Notably, increased transcription of imprinting loci *IGF2* and *H19* was observed at both day 3 and day 5 (Fig. S1C). Few selected *HOX* genes, except *HOXA1* and *HOXB1/2* (Fig. S1C), were activated, suggestive of cranial character of our NC progenitors.

**Figure 1.**
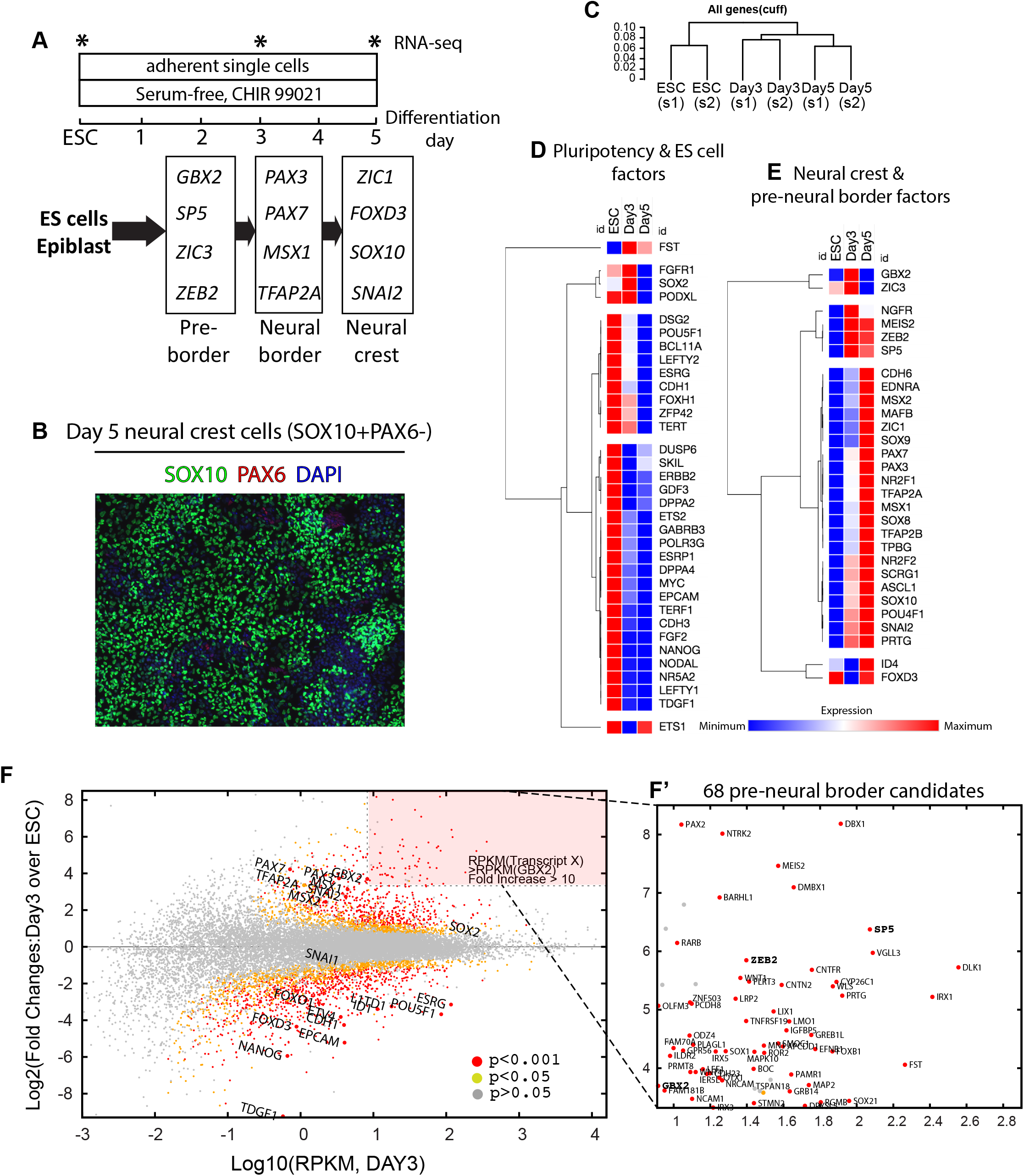
Identification of pre-border candidates by transcriptomic analysis. (A) Schematic diagram depicting the differentiation program for human NC induction. Asterisks indicate time points when samples were harvested for RNA-seq. (B) SOX10 and PAX6 immunostaining of day 5 NC cells. (C) Hierarchical clustering of all expressed genes from ESC, day 3 and day 5 transcriptomes. (D, E) K-means clustering of the RPKM values of transcripts mapped in the ESC, day 3 and day 5 transcriptomes that are functionally associated with pluripotency and self renewal of human ES cells (D), or neural plate border, pre-migratory neural crest and differentiating neural crest cells (E). Relative intensities were displayed with blue being lowest in expression level to red being highest. (F) Plot of all transcripts identified from RNA-seq with day 3 RPKM values plotted against day 3 fold change over day 0 (ES cells) (F’) Pink boxed-region is magnified on the right to display the 68 pB candidates transcripts. The 4 pre-border candidates identified in (Leung et al., 2016) are displayed in bold.

To systematically identify a complete pB gene set, we subjected day 3 transcriptome data to further analyses based on p values (<0.05), RPKM values (>8.2, expressing at a higher absolute transcript level than *GBX2*, a known pB gene), and fold changes (>10 folds over ES cells) of individual transcripts (Fig. 1F). Based on these criteria, 68 transcripts were identified (Fig. 1F’). These transcripts were expressed and induced at much higher level than neural border and NC transcripts. To examine the cell type specificity of these pB candidates, we performed RT-qPCR on 4 cell types: ES cells, pre-NC, prospective anterior neuroectoderm (NE) and non-neural ectoderm (NNE) (Fig. S1D). We found that 45.6% (n=31/68) and 19.1% (n=13/68) of pB candidates were “NC-specific” (Fig S1E) and “NC-enriched” (Fig S1F) respectively and were expressed at significantly higher levels in pre-NC cultures than those under ES cells, NE or NNE condition. For “NC-specific” pB candidates, no significant transcriptional induction was detected under NE and NNE conditions compared to ES (Fig S1E). These two categories together constitute 64.7% of all pB candidates. The rest of the candidates were either “NC-biased” (n=7/68, Fig S1G, induced under NC condition but no consistent increase in expression was observed compared NC versus NNE and NE), assuming a “broad ectoderm expression” pattern (n=7/68, Fig S1H, significant induction detected in all 3 ectoderm lineages), or “highly variable in expression or not specific to NC” (n=10/68, Fig S1I, no significant induction detected under NC condition). In conclusion, transcription of a majority of pB candidates was highly specific and enriched in early NC cultures.

### β-catenin dependent activation of early NC response genes

pB genes including *GBX2*, *ZEB2*, *ZIC3*, and *SP5* were known to be regulated by β-catenin (CTNNB1) signaling (Leung et al., 2016). As expected, β-catenin was the top predicted upstream regulator for differentially expressed genes found in our transcriptome (p=4.6e-27 and 1.36e-27 for day 3 and day 5 transcriptomes respectively, Fig. S2A). To test if pB candidates were β-catenin transcriptional targets, we examined the expression of 12 selected pB candidates, identified above as showing “NC-specific” induction (Fig S1E), in ES and differentiating cells that were transduced with either a *luciferase* shRNA or a *β-catenin* shRNA (Fig 2A). First we found that *β-catenin* shRNA downregulated *β-catenin* (*CTNNB1*) RNA expression at ES and day 1, but did not affect expression of pluripotency factor *POU5F1* (Fig. 2B). At day 1, a majority of pB candidates (7 out of 12) however were repressed by *β-catenin* shRNA (Fig. S2B). Notably, no change in pB candidate expression was detected in *β-catenin* shRNA cell line. By day 3, all 12 pB candidates were downregulated in *β-catenin* shRNA-transduced cultures (Fig. S2C). To confirm the expression dynamics of the rest of the pB candidates, we performed RNA-seq on cultures transduced with control *luciferase* or *β-catenin* shRNAs derived from ES cells, at day 1 and at day 3 (Fig. 2C, table S2). In line with our earlier time-course transcriptome analysis, 95.6% pB candidates (n=65/68) were induced in control *luciferase* shRNA cultures at day 3. Out of these candidates, 73.8% (n=48/65) displayed a 2-fold or more downregulation in *β-Catenin* shRNA cultures. A lower, yet significant, proportion of pB candidates (44.1%, n=30/68) were already induced in control day 1 cultures. Almost all of them (96.7%, n=29/30) were downregulated in *β-catenin* shRNA-transduced cultures.

**Figure 2.**
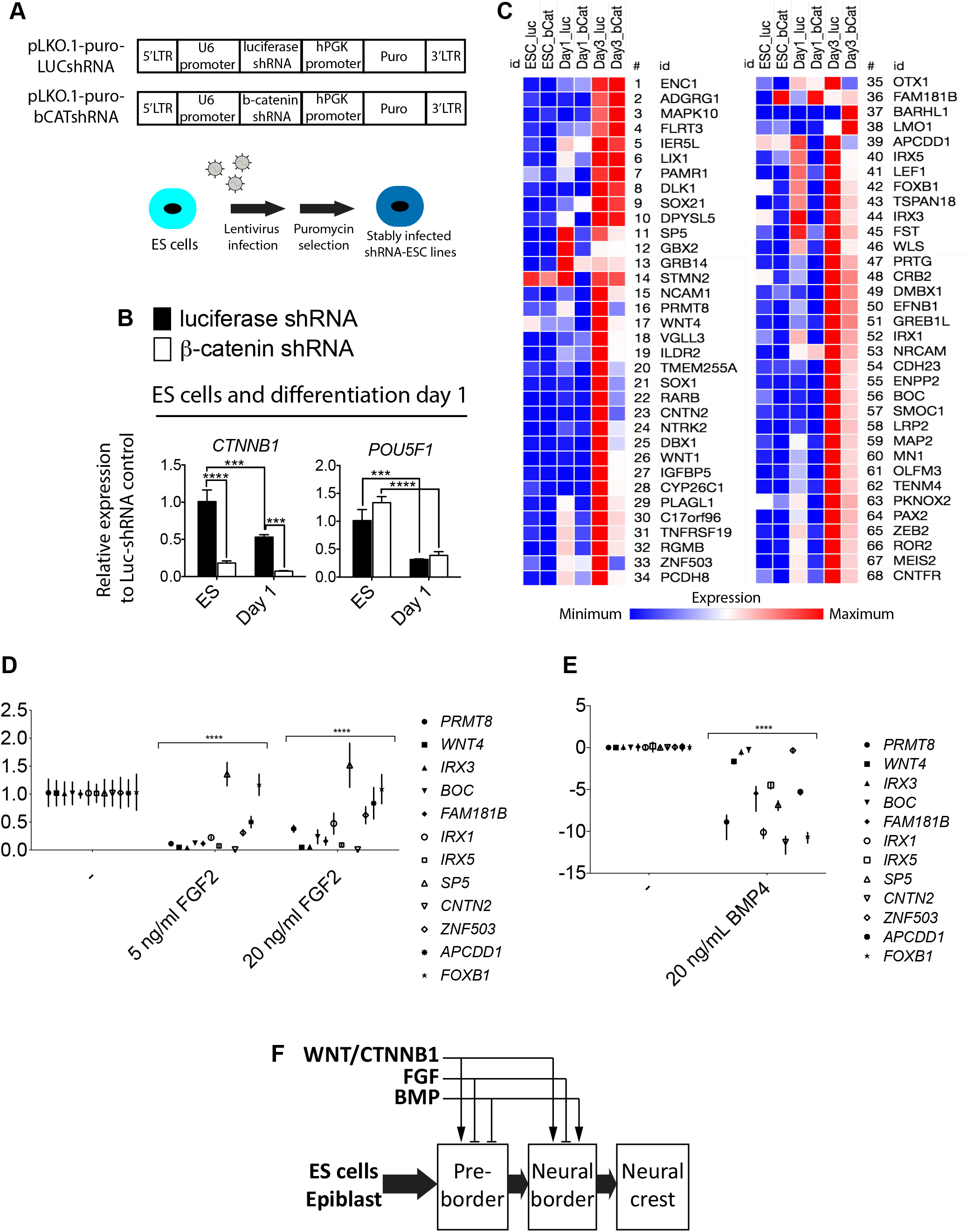
β-catenin positively regulates pB candidate gene induction. (A) Schematic diagram showing lentiviral constructs containing luciferase and β-catenin shRNAs and the procedures for generating stably infected shRNA expressing human ES cell lines. (B) qPCR data of *CTNNB1* and *POU5F1* comparing control (luciferase shRNA) against knockdown (*β-catenin* shRNA) in ES and day 1 cultures. (C) Heat map of the expression dynamics of the 68 pB candidates in control and knockdown cell lines derived from RNA-seq data of ES cell, day 1 and day 3 cells generated using Morpheus. (D) Differentiation day 3 cultures treated with or without FGF2 ligand at the indicated concentration. Dunnett’s multiple comparison tests were carried to calculate statistics between control group with ligand-treated groups. (E) Differentiation day 3 cultures treated with or without BMP4 ligand. Fisher’s LSD test was used. (F) A step-wise induction model for human NC and a gene regulatory network of human NC with signaling inputs from canonical WNT, FGF and BMP signaling.

FGF and BMP signaling are known to promote NC induction. To test if these signaling pathways contributed to the induction of pB candidates, we activated these pathways under NC condition using FGF2 (Fig. 2D) or BMP4 (Fig. 2E) ligand. Out of the 12 “NC-specific” pB candidates we tested, a majority of them were repressed to basal transcriptional level by either of these ligands suggesting that high FGF or BMP signaling did not favor pB candidate induction from ES cell cultures.

Altogether, these data establishes canonical WNT signaling is the main contributor among major signaling pathways for pB candidate induction (Fig. 2F) and β-catenin is a positive regulator for pB candidate induction during NC induction.

### Early NC response genes are Poised for Transcriptional Activation in Pluripotent Stem Cells

Expression of many pB candidates were robustly induced within 24 hours from the start of differentiation (Fig. S1C, E). It is possible that pluripotency factors expressed in ES cells might prime developmental genes such as pB candidates for which a rapid β-catenin activation was found.

By visual inspection of published ChIP-seq datasets for key pluripotency factors (NANOG, POU5F1, SOX2, BCL11A)(Gertz et al., 2013; Tsankov et al., 2015), β-catenin cofactors (NIPBL, LEF1, TCF4)(Estaras et al., 2015; Tsankov et al., 2015), and chromatin signature marks (Consortium, 2012; Rada-Iglesias et al., 2011), we identified putative regulatory elements close to pB candidate genes *GBX2* (Fig. S3A) and *FOXB1* (Fig. S3B). These 2 elements displayed strong binding of pluripotency factors and β-catenin co-factors, open chromatin configuration (ATAC-seq), presence of enhancer marks (EP300, H3K4me1), and β-catenin (CTNNB1) binding upon CHIR 99021(Funa et al., 2015) or WNT ligand (Estaras et al., 2015)-mediated ES cell differentiation. To identify other putative pB candidate regulatory elements, we tested the co-occupancy of pluripotency factors and β-catenin cofactors by performing pairwise comparisons of the ChIP-seq peak sets for pluripotency factors and β-catenin cofactors (Fig. S3C). We found significant overlaps for all pairwise comparisons (p<10^-4^) (Fig. S3C, Table S3). Among these comparisons, NANOG and NIPBL had the largest overlapping binding regions (n=37,481, Fig. S3E). Notably, these NANOG-NIPBL co-bound elements with characteristics of poised or inactive enhancers (Fig. S3F) were significantly associated with developmental processes for NC cells (NC differentiation/development, PNS development, abnormal craniofacial morphology) and gene expression in ectoderm (TS11/12/14 embryonic ectoderm) (Fig. S3G). These poised and inactive enhancers were also represented in high percentage (>50%) within the 199 NANOG-NIPBL co-bound elements associated with the 68 pB candidate loci (Fig. S3H). Among these 199 pB associated NANOG-NIPBL co-bound elements, a high percentage was also bound by β-catenin upon canonical WNT-mediated differentiation. Such elements were found at genomic location upstream (Fig 3A), downstream (Fig 3B), or within intronic sequences (Fig 3C) of pB candidate loci. To conclude, these *in silico* analysis supports the model that β-catenin signaling during NC induction unlocks poised or inactive enhancers for pB genes thereby leading to their increased transcription.

**Figure 3.**
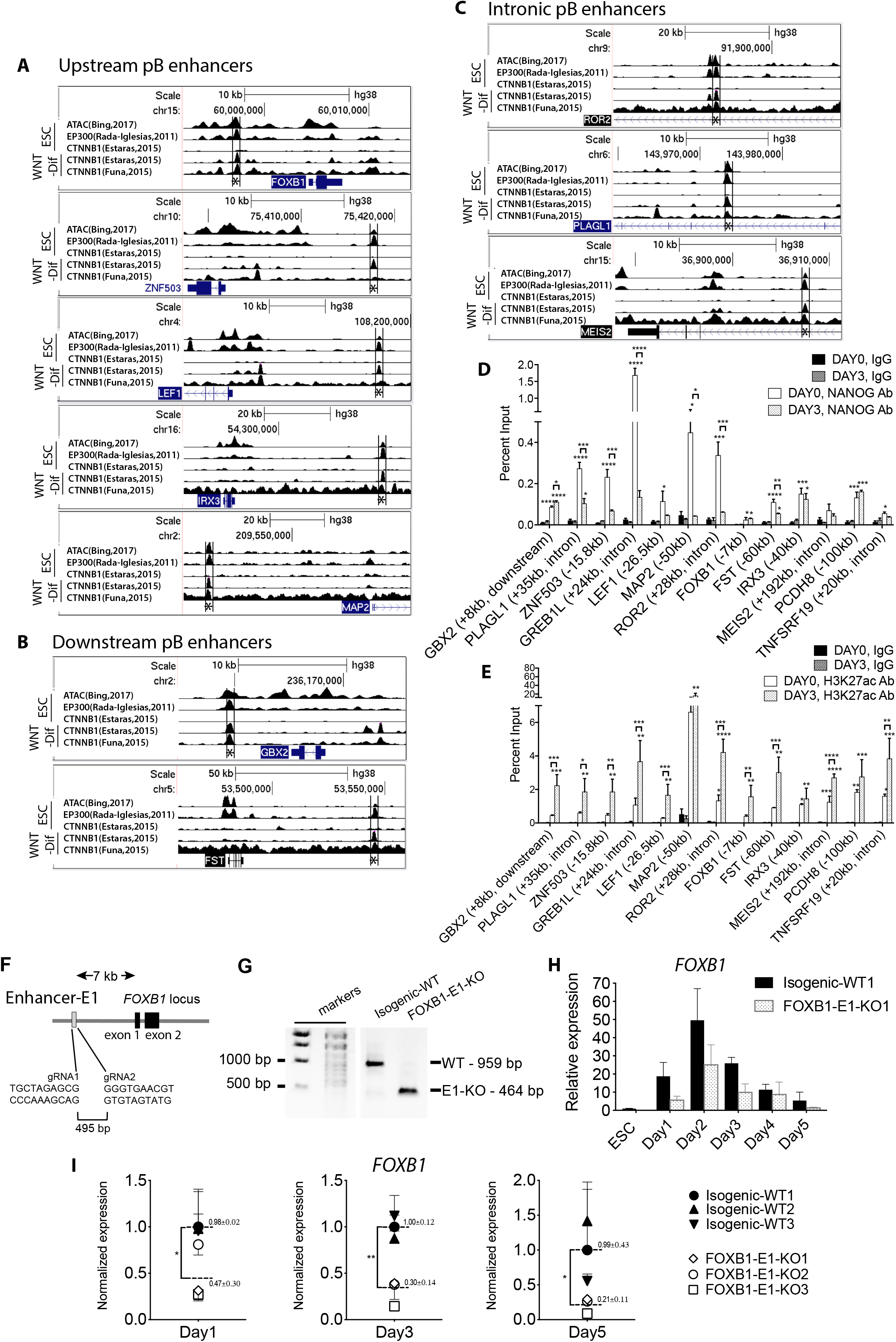
Functional characterization of pB gene regulatory elements. (A-C) Loci for pB candidate embedded with published ChIP-seq and ATAC-seq data sets in bigwig formats were displayed on UCSC browsers. ‘ESC stage’ indicates the sequencing data were derived from human ES cells. ‘WNT-Dif’ indicates the data were from human ES cells treated with CHIR 99021 or WNT3A ligand. qPCR measurements on ChIP DNA pulled down using anti-NANOG (D) and anti-H3K27ac (E) antibodies. Isotype-matched IgGs were used as controls. Asterisks directly above standard deviation bars represent comparisons between IgG and H3K27ac groups at their corresponding time points. One-way ANOVA tests were carried out with Bonferroni’s corrections for multiple testing. (F) CRISPR targeting strategy for FOXB1-E1 element. (G) PCR genotyping of isogenic and FOXB1-E1 targeted clones. Expected sizes of wildtype and knockout PCR products are shown on the right. (H) Day wise time course quantification of *FOXB1* expression during NC induction in isogenic-WT1 and FOXB1-E1-KO1 cell lines. Three culture samples were collected for each cell lines at each differentiation day. Unpaired t-tests with Holm-Sidak corrections for multiple testing were carried out on the indicated time points. (I) Normalized expression values and statistics at each time points were displayed.

To confirm the above *in silico* analysis that NANOG pre-occupied pB candidate regulatory elements and to visualize NANOG binding dynamics during NC induction, we performed NANOG CHIP-qPCR on ES and differentiation day 3 cells. We confirmed that 76.9% of all elements tested (n=10/13) have enriched NANOG binding at human ES cell stage. NANOG is known to prime developmental gene loci. During ES cell differentiation, NANOG is also known to repress NC gene transcription (Wang et al., 2012). While a smaller subset (n=6/13) still had significant NANOG binding at day 3, NANOG binding was much attenuated at day 3 as we detected an overall decreasing trend of NANOG binding in 53.8% of tested elements (n=7/13) at day 3 as compared to day 0 (Fig. 3D).

To demonstrate these enhancers were activated upon NC induction, ChIP-qPCR analysis the enhancer mark H3K27ac was carried on pB candidate regulatory elements in human ES cells (day 0) and differentiation day 3 cells. The data revealed that a majority of (76.9%, n=10/13) of the tested elements displayed increased accumulation of H3K27ac mark (Fig. 3C), demonstrating that these elements became more active during differentiation.

Lastly, to functionally test these pB enhancer elements, we used CRISPR approach to knock out one of them, the FOXB1-E1 (asterisk, Fig. 3B and 3E), a NANOG-NIPBL co-bound and β-catenin bound element that resides immediately upstream of the *FOXB1* gene. Deletion of this 495-bp E1 element (top panel, Fig. 3F) led to reduction of *FOXB1* transcription in differentiating cells in a day wise time course experiment (Fig. 3G). Further differentiation assay with additional knockout cell lines (FOXB1-E1-KO1, -KO2, -KO3) confirmed the time course experimental result in that removal of FOXB1-E1 element led to significantly reduced transcription of FOXB1 gene at differentiation day 1 (47%±30% of control level, p<0.01), day 3 (30%±14% of control level, p<0.001), and day 5 (21%±11% of control level, p<0.01) (Fig. 3H). The downregulation of *FOXB1* in differentiating cells strongly indicated that *E1* element acted as a functional enhancer for FOXB1 induction and its continued expression. These data together suggest that distinct regulatory elements primed by NANOG and NIPBL, and bound by β-catenin during early differentiation of human ES cells positively regulated pB candidate transcription.

### Pre-Border Candidates are expressed in chick epiblast prior to overt neural plate border formation

As a first approach to decipher the function of pB genes, we focus on FOXB1 which has a known role in lineage differentiation. Our RNA-seq data suggests that it is expressed at high level at differentiation day 3 during NC induction but not at ES cells or day 5 cultures (Fig 4A, S4A). This result is consistent with lineage tracing of *Foxb1* in murine embryos revealing its expression in NC precursors but not in mature NC cells or their derivatives (Zhao et al., 2007). By RT-PCR we confirmed that during human NC induction *FOXB1* was transiently induced and peaked at day 2 or day 3 similar to other documented pB genes (Leung et al., 2016), and its expression complemented two highly expressing FOX factors, *FOXD3* and *FOXH1,* which either rapidly downregulated upon differentiation (*FOXH1*) or assumed a bi-phasic expression pattern (*FOXD3*) (Fig. 3G, 4B). Whereas most other FOX factors *FOXA3*, *FOXI3*, *FOXN3*, and *FOXO4* showed decreased expression in day 3 compared to ESC, the only two other FOX factors that showed increased transcription at day 3 were *FOXOB3* and *FOXP4* but their fold increase in RPKM was limited to around 2 fold compared to more than 10 fold for *FOXB1*. In agreement to transcript analysis, FOXB1 protein was detected at day 3, but not in human ES cells (Fig. S4B). We reasoned that FOXB1 carried a unique function during the early differentiation window, when other FOX factors were not as highly expressed.

**Figure 4.**
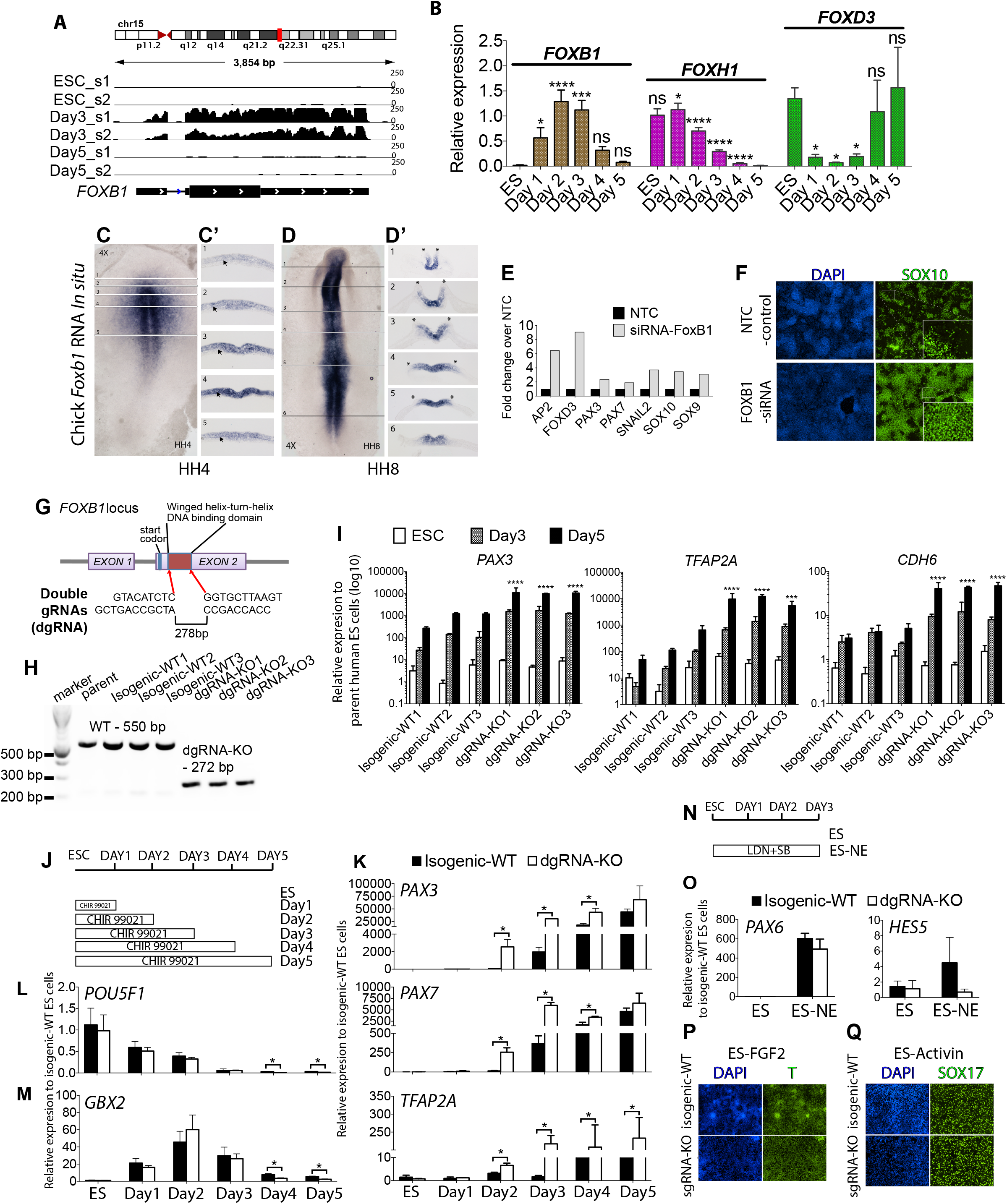
Phenotypic characterization of a human *FOXB1* loss-of-function cell model. (A) RNA-seq reads on human *FOXB1* locus converted to bigwig format and displayed on igv software. Sample labeling refers to Fig 1C. (B) Time course-qPCR analysis of selected FOX transcription factor transcripts during ES cell differentiation. (C-D) *Foxb1 in situ* hybridization in HH4 and HH8 stage chick embryos. Transverse sections are displayed to the right (C’ and D’). Arrows are pointing to the ectoderm cell layer. Asterisks indicate the position of the neural plate border. (E) qPCR data of NC factors on day 5 cultures treated with control or FOXB1 siRNA. A representative of 3 biological replicates is displayed. (F) DAPI staining and SOX10 immunostaining of day 5 cultures treated with control or FOXB1 siRNA. Magnified images for SOX10+ cells are displayed in the insets. (G) CRISPR targeting strategy for CRISPR-mediated mutations at the DNA binding domain sequence of *FOXB1* gene using two gRNAs. (H) PCR genotyping of isogenic and dgRNA targeted cell lines. (I) Time-course qPCR measurements of NC factors on three isogenic and dgRNA-KO cell lines. Two-way ANOVA was performed with Dunnett correction for multiple testing. (J) Differentiation and sample collection schematics for NC induction. (K) Time-course qPCR data of NC genes for isogenic wildtype and dgRNA-KO cell lines during NC induction. Multiple t-tests with Holm-Sidak corrections were performed. (L, M). *POU5F1* and *GBX2* time-course qPCR comparing isogenic control and *FOXB1* dgRNA-KO cell lines. Multiple t-tests with Holm-Sidak corrections were performed. (N) Differentiation and sample collection schematics for neurectoderm competency test. (O) Time-course qPCR data of neurectoderm genes during differentiation using a schematic displayed in panel N. Multiple t-tests with Holm-Sidak corrections were performed. Isogenic wildtype and sgRNA-KO ES cells were treated with FGF2 or Activin to promote mesoderm (P) and definitive endoderm (Q) induction as indicated by T and SOX17 immunostaining respectively.

Expression of *Foxb1* is first detected in gastrulating mouse embryos in the primitive streak and the surrounding epiblast (Labosky et al., 1997). In chick embryos, *Foxb1* was expressed at the same location at Hamburger-Hamilton (HH) stage 4 (Fig. 4C), posterior to the caudal boundary of the anterior neural marker *Ganf* (Fernandez-Garre et al., 2002), and just before *Pax7* induction at the prospective neural border at HH4+ (Basch et al., 2006). *In situ* hybridization for other pB candidates with chicken homologs including *Cntn2*, *Prmt8*, *Nrcam*, *Plagl1*, *Stmn2*, *Enpp2*, and *Enc1* showed similar expression patterns as *Foxb1* at HH stage 4 (n=7, Fig. S4C). Notably, *Foxb1* robust expression is restricted to the ectoderm/epiblast layer (arrows, Fig. 4C’). Upon development, *Foxb1* expression progressively restricted medially as neural plate developed (Fig. 4D), and was visibly extinct from the neural border at HH8 (asterisks, Fig 4D’). The broad expression domain of *Foxb1* in early epiblast cells could represent a competent zone for NC development, and hinted at a function for Foxb1 in prospective NC populations prior to the formation of the neural border.

### Loss of FOXB1 De-Repressed Neural Crest Development

To interrogate the function of FOXB1 in human ES cell differentiation, we attempted to reduce expression level of FOXB1 induced by administration of a *FOXB1*-siRNA. Surprisingly, FOXB1-siRNA resulted in upregulation of NC marker expression at day 5 as indicated by increased transcription of NC markers including *SOX10* (Fig 4E) and proportion of SOX10^+^ cells (Fig. 4F). We then took advantage of CRISPR/CAS9 technologies. To this end, we removed the sequence encoding the Forkhead DNA-binding domain of FOXB1 using a pair of guide RNAs (dgRNA) (Fig. 4G). PCR genotyping and Sanger sequencing revealed specific deletion of the 278-bp genomic region containing the sequence encoding the FOXB1 DNA-binding domain (Fig. 4H and data not shown). Analysis with Western blot confirmed the loss of FOXB1 protein in differentiating *FOXB1-KO* cell lines (Fig. S4B). We then assessed possible effects during NC formation in KO lines and isogenic controls. Cultures from the parent human ES cell line, 3 isogenic and 3 KO cell lines were processed for RT-qPCR analysis at ES cell stage, day 3, and day 5. Confirming siRNA data, expression of neural border and NC genes *PAX3*, *TFAP2A*, and *CHD6* were significantly upregulated at day 5 in KO over control lines (Fig. 4I). Increases in expression for these genes however were already observed at day 3, suggesting that loss of FOXB1 affected the early phase of NC induction. Consistent with qPCR data, immunostaining of dgRNA-KO cells at day 5 also revealed increased SOX10+ cells (data not shown). To confirm the dgRNA experiment, we designed another experiment that targeted the translational start site of FOXB1 using a single guide RNA (sgRNA) (Fig. S4D). We established a knockout (sgRNA-KO) and a heterozygote (sgRNA-Het) cell line for analysis. We characterized the mutations in these 2 cell lines by TA-cloning (Fig. S4E) and verified the absence of FOXB1 protein in sgRNA-KO line by western blotting (Fig. S4F). To provide a broader perspective of the influence of FOXB1 in NC development we performed RNA-seq on these samples. Principal component analysis of sgRNA-KO and sgRNA-Het transcriptome data of differentiating cells collected at day 3 and day 5 revealed clear segregation of the isogenic control, sgRNA-Het and sgRNA-KO cell lines according to their differentiation time points by PC1, as well as tight correlations according to their genotypes at PC2, revealing a potential dosage effect of FOXB1 function (Fig. S4G). Similar to dgRNA-KO cell lines, we found that neural border and NC transcripts were upregulated in sgRNA-Het and sgRNA-KO at both day 3 and day 5 (asterisks, Fig. S4H, table S4). A few pB candidates were also upregulated (*DMBX1*, *OLFM3*, *PAX2, PKNOX2*) or downregulated (*NRTK2*, *SOX21*, *VGLL3*) indicating that FOXB1 might feedback control selected pB genes (double asterisks, Fig. S4H). Intriguingly, we also observed activation of imprinting loci (*H19*, *IGF2*, *XIST*)(Fig. S4I), PRC2-regulated loci (*HOXA* and *HOXB* clusters, as well as the neighboring homeodomain gene *EVX1*) (Fig. S4J and data not shown), and loci related to increased cell proliferation (*EGFR, MYC*, and *S100A11*) (Fig. S4K).

To investigate more precisely when does FOXB1 affects NC induction, we performed time-series analyses (Fig 4J). We found that dgRNA-KO cells underwent accelerated NC induction, as revealed by a 24-hour earlier induction of neural border genes *PAX3*, *PAX7*, and *TFAP2A* (Fig. 4K). Instead of an accelerated loss of pluripotency marker expression as would be expected, expression dynamics for pluripotency gene *POU5F1* appear similar in isogenic control and dgRNA-KO cells (Fig. 4L). Similarly, expression of another pB gene *GBX2*, seem normal from day 1 to 3 in control and dgRNA-KO cells (Fig. 4M). These results suggest that FOXB1 may regulate the temporal acquisition of border specifiers and later NC markers, independently of POU5F1, and GBX2. Then we interrogated if FOXB1 depletion modulates other fates. To this end we deployed a BMP and Activin inhibitor cocktail on ES cell cultures to promote neurectoderm fate (Fig. 4N). Control and dgRNA-KO FOXB1 cells appear to attain neuroectoderm precursor status and displayed no significant expression differences in neurectoderm markers *PAX6* and *HES5* (Fig. 4O), even though *HES5* dgRNA-KO displayed a reduced transcription trend.

As indicated by comparable expression, loss of *FOXB1* also did not alter the ability of dgRNA-KO cells line or sgRNA-KO cells to form mesoderm (T, Fig. 4P) or definitive endoderm (SOX17, Fig. 4Q)(ES and ES-de, *EOMES*, *CER1*, and *SOX17*, Fig. S4L, M) from ES cell stage, suggesting that FOXB1 effect on ES cell differentiation was specific to NC lineage. These data altogether demonstrated that loss of FOXB1 promoted the acquisition of NC.

### FOXB1 Directly Regulated the Transcription of Key Neurogenic and Neural Crest Loci

To interrogate the molecular mechanism by which FOXB1 acted on the NC differentiation program, we examined the genome-binding pattern of FOXB1. We performed ChIP-seq of day 3 cultures when FOXB1 protein could be detected using a specific anti-FOXB1 antibody (Fig. S4B, S4F). Peak calling against input sample uncovered 3348 and 4388 high-confidence peaks for 2 wildtype ES cell samples (FDR<0.1%, Day3WT1 and Day3WT2, Fig. 5A) that had significant overlaps with each other (enrichment fold log_2_=9.3, p<10^-5^, Fig 5A’), but only ∼10 peaks each for 2 dgRNA-KO samples (FDR<0.1%, Day3KO1 and Day3KO2, Fig. 5A). The 2158 day-3 common peaks had enrichment of GC-rich repeats and forkhead transcription factor-binding motif (data not shown and Fig. 5A”), and were specifically bound by FOXB1 as shown by heat maps of aggregating ChIP-seq signals for wild type, dgRNA-KO, and input samples (Fig. 5B). Specificity of FOXB1 binding was additionally verified by ChIP-qPCR assay against IgG and dgRNA-KO controls at selected FOXB1 bound regions (n=13, Fig. S5A).

**Figure 5.**
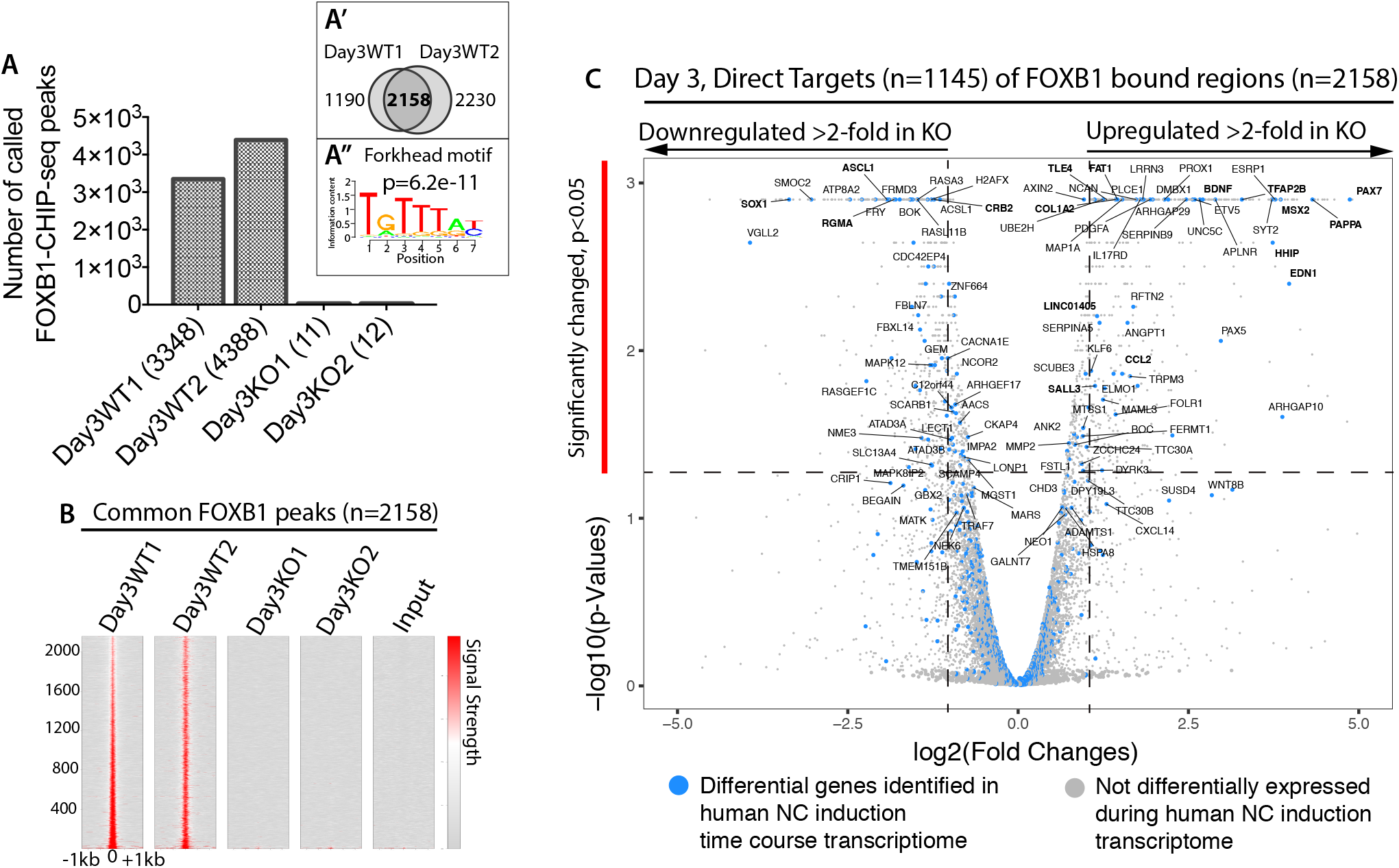
Global genomic binding analysis of FOXB1 protein during human neural crest induction. (A) Peak numbers called from FOXB1 ChIP-seq on 2 wildtype and 2 dgRNA-KO samples. (A’) Venn diagram showing overlapping elements for the 2 wildtype ChIP-seq samples. (A”) Motif analysis of 2158 FOXB1-bound elements. (B) Heat map of signals centered on the 2158 FOXB1-bound elements for 2 wildtype, 2 knockout and an input samples. (C) Volcano plot of all FOXB1 targets within a 50kb window from TSS. Purple dots indicate genes that are differentially expressed in our earlier ES, day 3 and day 5 transcriptome (Fig 1). Bolded genes are key candidates for neural and NC development.

Volcano plot analysis of direct FOXB1 targets combining our above FOXB1 sgRNA-KO RNA-seq data and the FOXB1 CHIP-seq data set revealed 3 classes of gene targets (Fig. 5C, Table S5). The first class included neural progenitor genes that were downregulated (>2-fold in RPKM) in the FOXB1 sgRNA-KO day 3 transcriptome such as *ASCL1*, *CRB2*, *RGMA* and *SOX1* (bolded, Fig. 5C). Of interest, *CRB2* and *SOX1* were also identified as pB candidates (Fig. 1F’). The second class included FOXB1 direct targets whose transcription were not affected in FOXB1 sgRNA-KO cells at day 3. Among them, there were genes whose expression was normally increased in our differentiating cultures such as *IGF1R* and *JAG1* (Fig. S1B), and several pB candidates *BOC*, *CNTFR*, *GBX2*, *GREB1L, MAP2*, *MEIS2*, *OTX1* and *ZNF503* (Fig. 1F’). Neural progenitor and differentiation genes (*FEZF2*, *ISL1*, *MSI1*, *POU6F1*), placodal ectoderm regulator (*SIX1*), and general DNA methylation machinery components (*DNMT1* and *DNMT3A*) were among FOXB1 targets whose expression was not affected at day 3 (Table S4). Lastly, the third class included upregulated genes (>2-fold in RPKM) such as *BDNF*, *CCL2, COL1A2*, *DMBX1, HHIP, LINC01405, MSX2*, *PAPPA*, *PAX7*, *FAT1, SALL3*, *TFAP2B,* and *TLE4* (bolded, Fig. 5C). Among the upregulated gene loci, some of them were known NC genes such as *EDN1*, *MSX2*, *PAX7*, and *TFAP2B*, whereas others were pB candidates (*DMBX1*, Fig 1F’) or normally upregulated during NC induction including *CCL2*, *COL1A2*, *FAT1*, *HHIP*, *PAPPA*, and *TLE4* (Fig. S1B and table S1).

By visualizing FOXB1-binding sites within target gene loci, we found that for neural progenitor gene loci, FOXB1-binding sites could be found upstream of *ASCL1*, *SOX1*, and *MSI1* genes (asterisks, Fig. 6A-C). Among the FOXB1-bound elements in the upregulated NC gene loci, some were intronic or located within gene bodies (*PAPPA, PAX7*, *SALL3*, *TLE4*, *FAT1*, *HHIP*, and *TFAP2B* loci)(asterisks, Fig. 6D, E, S6A); others were located either upstream (*BDNF*, *CCL2*, *COL1A2*, and *EDN1* loci) (asterisks, Fig. 6F, S6B) or downstream (asterisks, *MSX2* locus) (Fig. 6G).

**Figure 6.**
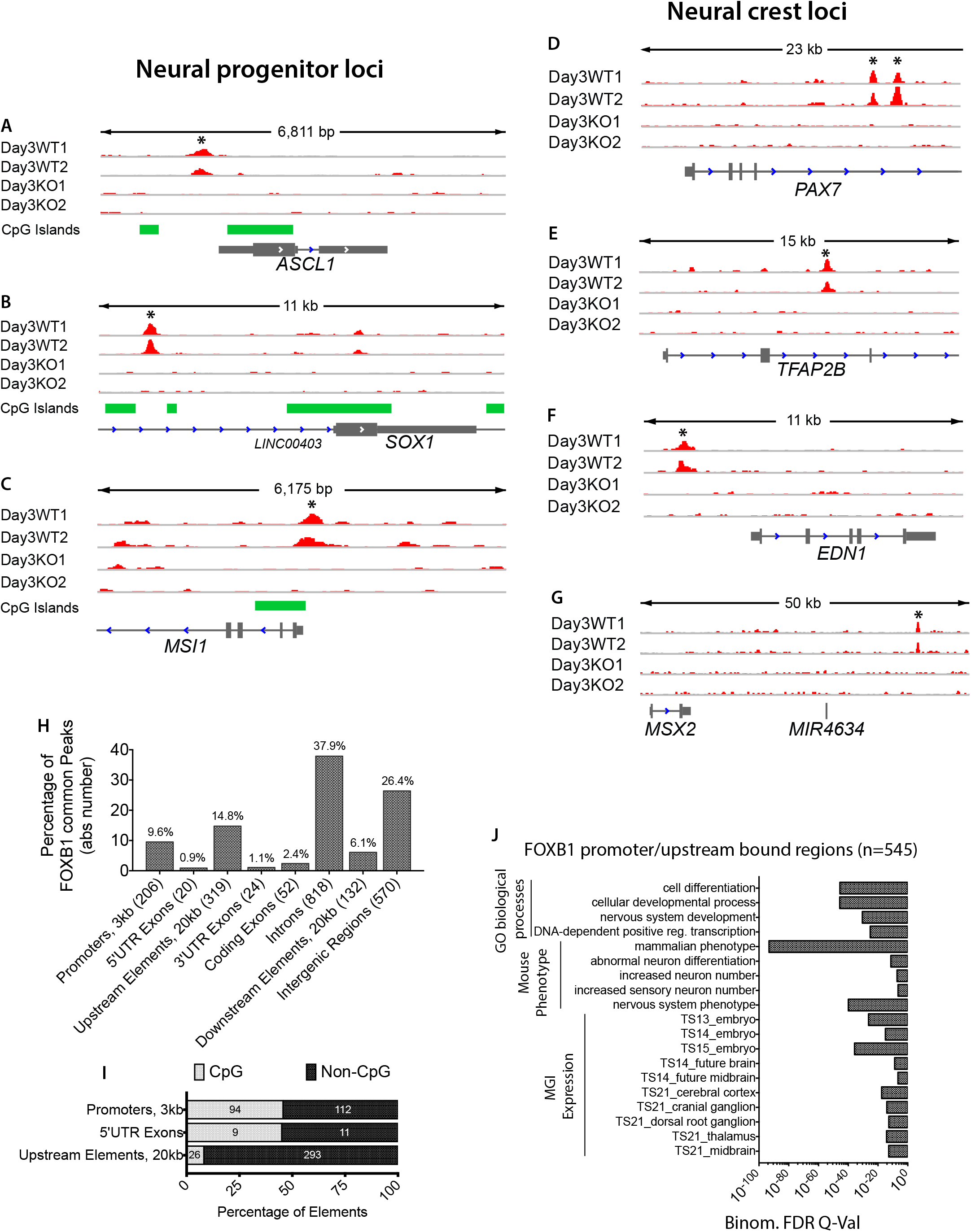
FOXB1 directly regulates target neural and neural crest genes. IGV tracks displaying ChIP-seq signals for the 2 wildtype, 2 knockout samples, and an optional CpG annotation at neuronal progenitor (A-C) and neural crest (D-G) gene loci. Asterisks indicate location of called peaks. (H) Genome annotation of the 2158 common FOXB1 peak regions using NCBI Ref. Seq. build hg19. Priority was set as follows: Promoters, 5’UTR exons, upstream sequences, 3’UTR exons, coding exons, introns, downstream sequences and intergenic regions. Actual peak numbers are shown in brackets. (I) The 545 selected FOXB1-bound elements, contained within upstream elements, promoters, and 5’ UTR exons, were further classified according to the presence or absence of CpG islands. (J) GREAT gene ontogeny analysis of the 545 selected FOXB1 peak set using a 20 kb genomic window.

Genome annotation revealed a significant proportion of intronic FOXB1-bound elements and an enrichment of FOXB1-bound elements locating within promoters (9.6%) and upstream regions (14.8%) (Fig. 6H). We found that many FOXB1-bound elements within 5’ UTR exons and promoters were either close to (Fig. 6A-B) or contained (Fig. 6C) CpG islands. We performed gene ontogeny analysis on a shortlisted FOXB1-bound element set containing only 5’ UTR exons, promoters and upstream elements (n=545). We found significant enrichment of FOXB1 direct targets with gene sets correlated to mammalian-specific phenotype, neuronal differentiation, and developmental processes (Theiler stages 13 to 21) at peripheral (NC-derived) and central (neuroectoderm-derived) nervous system structures (Fig. 6J) consistent with the model that FOXB1 directly regulate human NC cell differentiation. In conclusion, our CHIP-seq data confirmed that FOXB1 directly regulate NC gene induction by directly binding to NC and related gene loci.

### Loss of FOXB1 Function Leads to Long-term Suppression of Autonomic Neuronal Gene Expression

Transient expression of a transcription factor can confer developmental competence to differentiating cells to generate subsequent lineages as has been demonstrated for other transcription factors such as Neurod1 (Pataskar et al., 2016) and Foxa1/2 (Wang et al., 2015). The binding of FOXB1 in day 3 pre-NC cells to neural progenitor gene loci could therefore carry such developmental function. One of these neural progenitor loci, ASCL1, its gene product is a master regulator for neurogenic potential as well as autonomic neuron differentiation (Lo et al., 1998; Oh et al., 2016). NC contributes to mesenchymal progenitors and peripheral neurons including sensory and autonomic neurons. We therefore speculated that removal of FOXB1, which directly regulated *ASCL1*, would have a far-reaching effect on the differentiation program of human NC cells towards autonomic lineages. While we also hypothesize that effect on *ASCL1* expression or other direct targets may affect the balance of mesenchymal and other neurogenic potentials in FOXB1 KO cell lines.

To test these hypotheses, we performed differentiation assay of day 5 isogenic wildtype and dgRNA-KO cells (Fig. 7A) using a tri-inhibitor cocktail (CHIR 99021, SU5402 and DAPT) to promote peripheral neuron (PN) formation (Chambers et al., 2012; Leung et al., 2016), or FGF2 ligand to promote mesenchymal progenitor (MP) formation (Fig. S7A-B). We observed that autonomic neuronal markers *ASCL1* (Fig. 7B), *PHOX2A* (Fig. 7C), and *TH* (Fig. 7D), sensory neuron marker *POU4F1* (Fig. 7E) and general neuronal marker *TUBB3* (Fig. 7F) were induced in day 11 isogenic control cells under PN, but not under MP, condition, confirming that PN condition promotes both sensory and autonomic neuron generation. Notably, *ASCL1* (Fig. 7B) and its downstream targets in autonomic neuron development, the homeodomain transcription factor (*PHOX2A*)(Fig. 7C), and the enzyme synthesizing the neurotransmitter noradrenaline (*TH*) (Fig. 7D), were either not induced to the same extent as isogenic control cells or not induced at all in dgRNA-KO under PN differentiation condition. Expression of general or sensory peripheral neuron markers, such as *POU4F1* (Fig. 7E) and *TUBB3* (Fig. 7F), however, were not changed in dgRNA-KO cells. Intriguingly, we observed that mesenchymal markers *TWIST1* (Fig.7G) and *NT5E* (Fig.7H) were upregulated in day 5 and/or day 11-MP dgRNA-KO cells upon MP induction.

**Figure 7.**
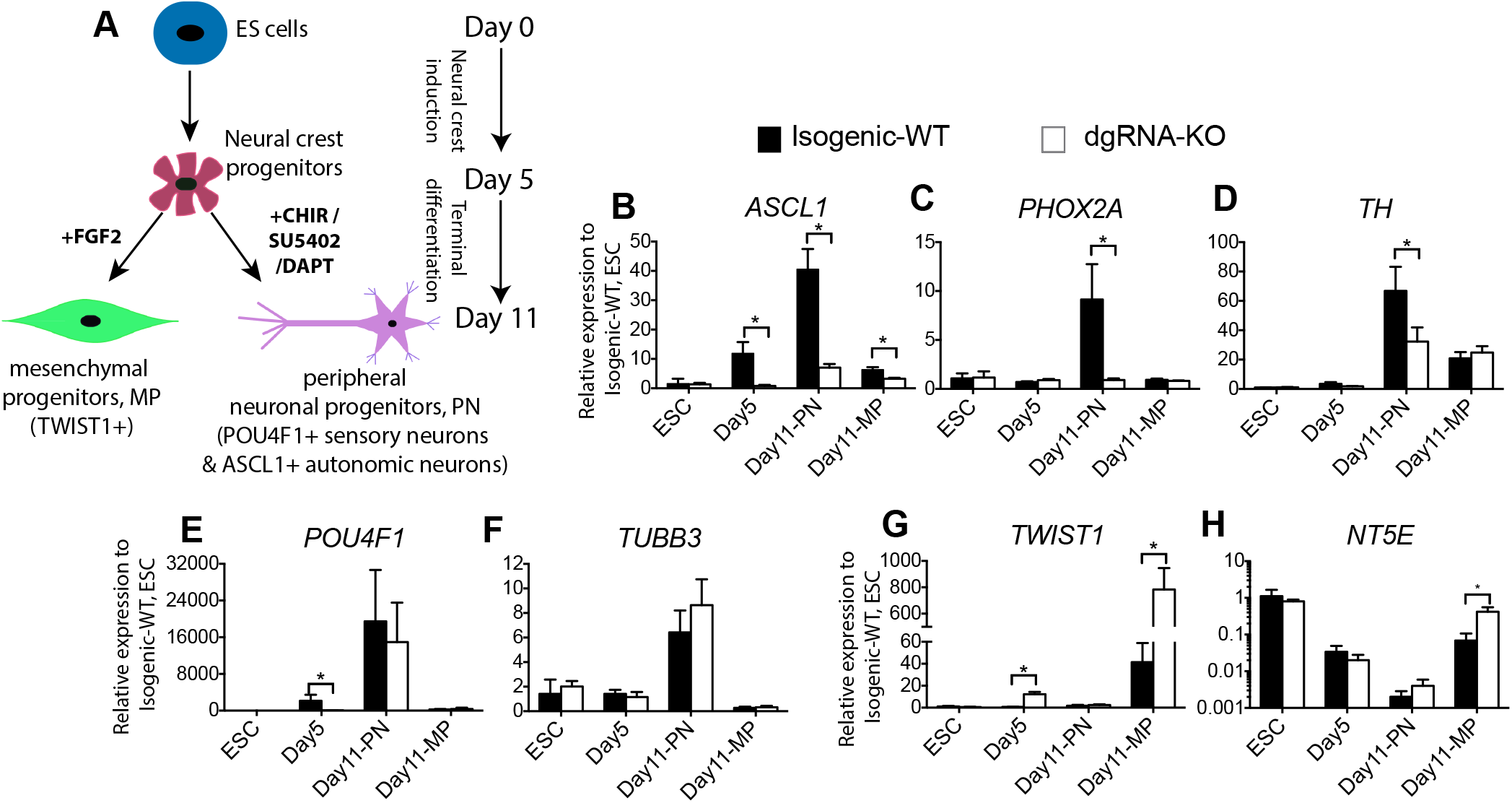
*FOXB1* regulates autonomic neuron differentiation potential in NC cells. (A) Terminal differentiation schematic for NC cells. qPCR analysis of autonomic neuronal (B-D), sensory and general neuronal (E-F) and mesenchymal progenitor (G-H) markers on ES, day 5 and differentiated day 11 isogenic control and dgRNA-KO cells.

These data suggested that loss of FOXB1 function promoted expression of mesenchymal genes but had a long-lasting effect on *ASCL1* gene transcription and might therefore affect the ability of mature NC to generate autonomic neuronal progenitors as indicated by inability of FOXB1 KO to upregulate other autonomic markers.

## DISCUSSION

Identification and characterization of novel cell intermediates, especially those that exist transiently in early embryos, is a challenging process. Here, we present a pipeline for such research, which utilizes a range of techniques including time-course transcriptomic analysis to identify key transcripts, signaling pathway perturbations to characterize signal dependency, knockout of enhancers to demonstrate regulatory functions, gene targeting of candidate factors and differentiation assays for phenotypic characterization, and genome-wide chromatin binding data to pinpoint genomic occupancy and to infer direct regulations and genetic networks. This pipeline could be applied to studies of other cell types in stem cell and regenerative biology to identify novel cellular states and intermediates. Our data here supports the existence of pB state, however more definitive answers awaits further evidence, perhaps from the use of approaches such as single cell RNA-sequencing and lineage tracing experiments would provide more definitive proof. In this study, we also provide valuable resources and genomic data sets for FOXB1 binding, β-catenin immediate targets during ES cell differentiation, putative novel enhancer location for pB candidates/β-catenin and pluripotent factor regulated loci, as well as transcriptome of NC induction for data mining and for generation of further testable hypotheses.

Our pioneer discovery of pB as a cellular intermediate during NC induction (Leung et al., 2016) is gaining momentum and increasing attention in recent years in embryo and reprogramming studies. For instance, a recent work describes a novel population of ‘neural border stem cells’ reprogrammed from blood cells that bear potential to generate central nervous system and NC derivatives (Thier et al., 2019). In another study, Trevers et al., identifies embryonic ‘pB’ cells from pre-gastrula avian embryos that bear resembling molecular signatures and differentiation potentials to our *in vitro*-derived pB cells (Trevers et al., 2018). Our current manuscript serves as a follow-up study for our previous work and others, and provides the molecular framework to derive pB-like cells from embryos and other differentiated cell types for developmental, stem cell, and therapeutic studies.

While inhibition of bone morphogenetic protein (BMP) signaling abolished human NC induction and neural border gene expression, expression of pB genes was preserved (Leung et al., 2016). In our expanded list of 68 pB candidates, we also found that in almost all candidates tested including FOXB1, induction of these genes were immune to inhibition of BMP signaling (data not shown). Such difference in responses to BMP signaling hints at a potential strategic function of pB candidates in NC development.

Our data identifies FOXB1 as a key factor limiting NC induction. Here, we show that FOXB1 is expressed transiently in human pre-NC cells before the specification of NC. We additionally show that *Foxb1* expresses in chick embryos prior to the appearance of the neural plate border. Lineage tracing in mouse embryos supports the notion that Foxb1 is expressed in pre-NC cells and there is no evidence to suggest that it is present in mature NC cells or their derivatives.

Our findings from FOXB1 ChIP-seq and the phenotypic analyses of FOXB1 KO cells suggest a broader function of FOXB1 in directly regulating NC development. FOXB1 function however is distinct from FOXD3, a pluripotent stem cell and NC factor, whose expression complements FOXB1 during NC induction, and acts as an activator for NC development. The fact that the neural border gene *PAX7* is directly regulated by FOXB1 strongly indicates that FOXB1 acts upstream and represses the development of the neural border stage. Sequence conservation analysis by MultiZ (Emera et al., 2016) reveals that our FOXB1-bound elements are less evolutionary-conserved. More than 5% of these elements are originated from the human clade (compared to <1% in control elements) and up to 25% are primate or younger sequences (data not shown). We postulate that evolution of these young elements could regulate NC developmental processes, thereby shaping species-specific features during human and primate evolution. Indeed, we found strong primate-specific FOXB1 bound elements within key NC loci such as *MSX2* (+41K), *EDN1* (−131K), and *NR2F1* (−91K), which could potentially contribute to the discrepancy between our data and murine embryo data that shows no alteration of border/NC markers such as *Pax3* and *Msx1* in *FOXB1* mutant mice (Labosky et al., 1997).

It is assumed that NC cells retain neurogenic potential from neurectoderm progenitors, which NC was believed to derive from. But, here and in our previous work (Leung et al., 2016), we have shown that NC and neural lineages segregated from each other before the appearance of definitive neuroectoderm. NC cells therefore have to establish its neurogenic potential in a mechanism either shared with or independent of pre-neuroectoderm progenitors. We found that FOXB1 KO cells acquired a heightened differentiation potential to generate mesenchymal progenitors. This was accompanied by a deficiency in their ability to specify autonomic neuronal fate. Such defect in differentiation potential could be due to a change of axial identity in FOXB1 KO cells (which show an upregulation of anterior *HOX* genes), as migrating NC populations from different axial identity of the rhombomeres are known to differentially contribute to autonomic neuronal lineages (Lumb et al., 2014). Our CHIP-seq data however shows that FOXB1 is directly activating and maintaining the competency of NC cells to express *ASCL1*, an autonomic neuron master regulator (Lo et al., 1998). FOXB1 may act via maintaining the open chromatin structures at *ASCL1* proximal promoter and/or other unknown epigenetic mechanisms to maintain NC cell competency to induce *ASCL1* expression. On the other hand, it could act as a classic transcription factor via recruitment of co-activators or co-repressors to activate or repress target gene expression similar to other FOX family proteins (Lalmansingh et al., 2012), or specific domains within the FOX factor could specify these distinct transcriptional activities as have previously shown for other FOX factors (Neilson et al., 2012).

Interestingly, we also find that a significant portion of our FOXB1 peaks (>30%) are occupied by the neuron subtype-specific transcription factor and an autism susceptible gene FOXP1 in human neural progenitor cell lines (Araujo et al., 2015)(data not shown) suggesting that FOXB1 may carry unknown functions in regulating latter neural development.

The early requirement of WNT/β-catenin signaling for human NC induction (Leung et al., 2016; Menendez et al., 2011; Mica et al., 2013) has been reported in vertebrate model organisms such as birds, amphibians and fish (Basch et al., 2006; Chang and Hemmati-Brivanlou, 1998; Garcia-Castro et al., 2002; Patthey et al., 2009; Saint-Jeannet et al., 1997)(Prasad et al., submitted). Our data supports a model in which WNT/β-catenin signaling directly activates a panel of pB genes including FOXB1 and other factors such as GBX2 and SP5 that in turn regulate the specification, lineage competency, and differentiation timing of differentiating ES cells to acquire NC fate. Previous works have demonstrated pre-broder genes GBX2 and SP5 acting upstream of neural border genes to induce the latter expression (Li et al., 2009; Park et al., 2013). We propose that pro-neural crest factors such as GBX2 and SP5 could act parallel to FOXB1 to influence NC fate choice.

This work further suggest that the proposed transient pB cell stage has unique transcriptional, epigenetic and functional features, different from the pluripotent cell state from which it emerges, and from later stages NC progenitors found at the neural plate border and neural folds. We have confirmed early expression for several pB gene candidates prior to expression of neural plate border genes in chick embryos, and demonstrated in our human system their early expression to be canonical WNT dependent and insensitive to BMP inhibition. Focusing on FOXB1 as a pB gene example, we showed that it directly regulates NC loci, temporally controls progression of differentiating cells towards more advanced NC state, and carries a remarkable impact in terminal differentiation driven by its early transient expression. These findings lend strong support to our proposed pB state and its significance to NC ontogeny.

## Supporting information

Supplemental text including methods and materials

## ACKNOWLEDGEMENTS

We thank Dr. Caihong Qiu, Yinghong Ma, and Jason Thomsen at the Yale Stem Cell Core for providing access to core facility and human pluripotent stem cell lines, and Dr. Kaya Bilguvar, Christopher Castaldi, and Bryan Pasqualucci at the Yale Center for Genome Analysis (YCGA) and Dr. Mei Zhang at the Genomics and Bioinformatics Core for high throughput sequencing services. We thank Dr. James Noonan for his critical advice on the design of the FOXB1 ChIP-seq experiment. We also thank Dr. Giuseppe Amatulli and Dr. Dejian Zhan for providing assistance on bioinformatics analysis. All public data sets were downloaded from Cistrome database. This work is supported by funding from Connecticut Innovations and NIH R01DE017914.

## AUTHOR CONTRIBUTIONS

Conceptualization: A.W.L., A.Z.X., M.I.G.C.; Methodology: A.W.L., C.B.; Software: F.L-G; Writing - Original Draft: A.W.L.; Writing – Review & Editing: A.W.L., A.Z.X., M.I.G.C; Formal analysis: A.W.L., F.L-G; Investigation: A.W.L., B.C., M.S.P., J.C.; Visualization: A.W.L., F.L-G, K.L.; Funding acquisition: A.Z.X., M.I.G.C.

