## Supplemental figures 1 to 7 for "Pre-Border Gene Foxb1 Regulates the Differentiation Timing and Autonomic Neuronal Potential of Human Neural Crest Cells"

SUPPLEMENTAL FIG S1 RELATED TO DATA IN FIGURE 1

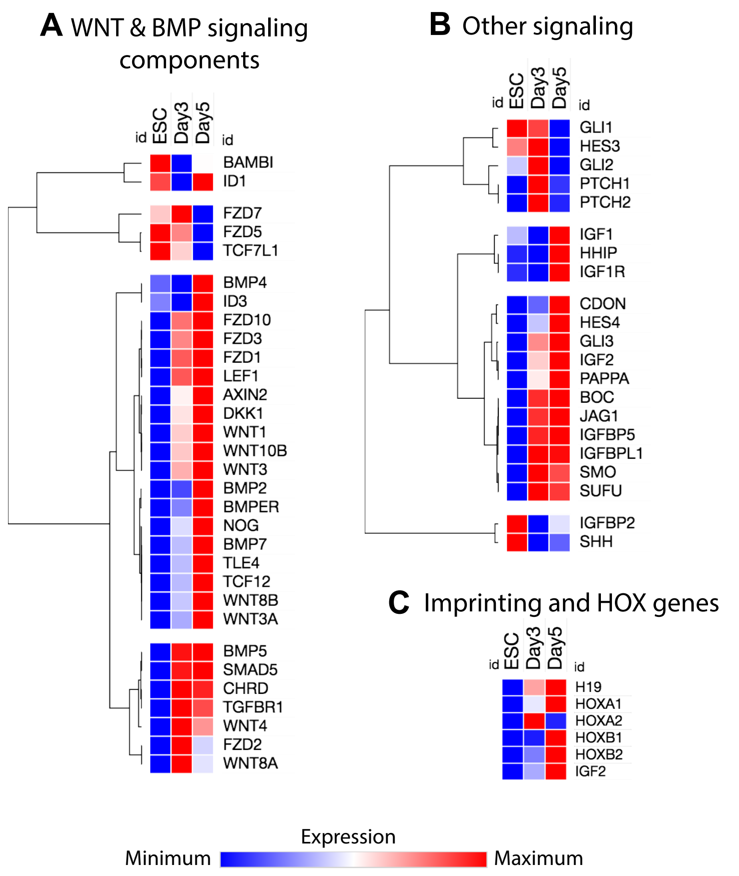

SUPPLEMENTAL FIG S1 RELATED TO DATA IN FIGURE 1

D

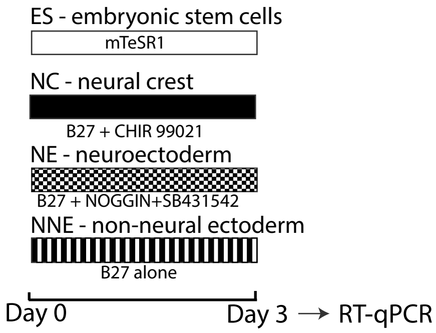


### E Neural crest specific (N=31)

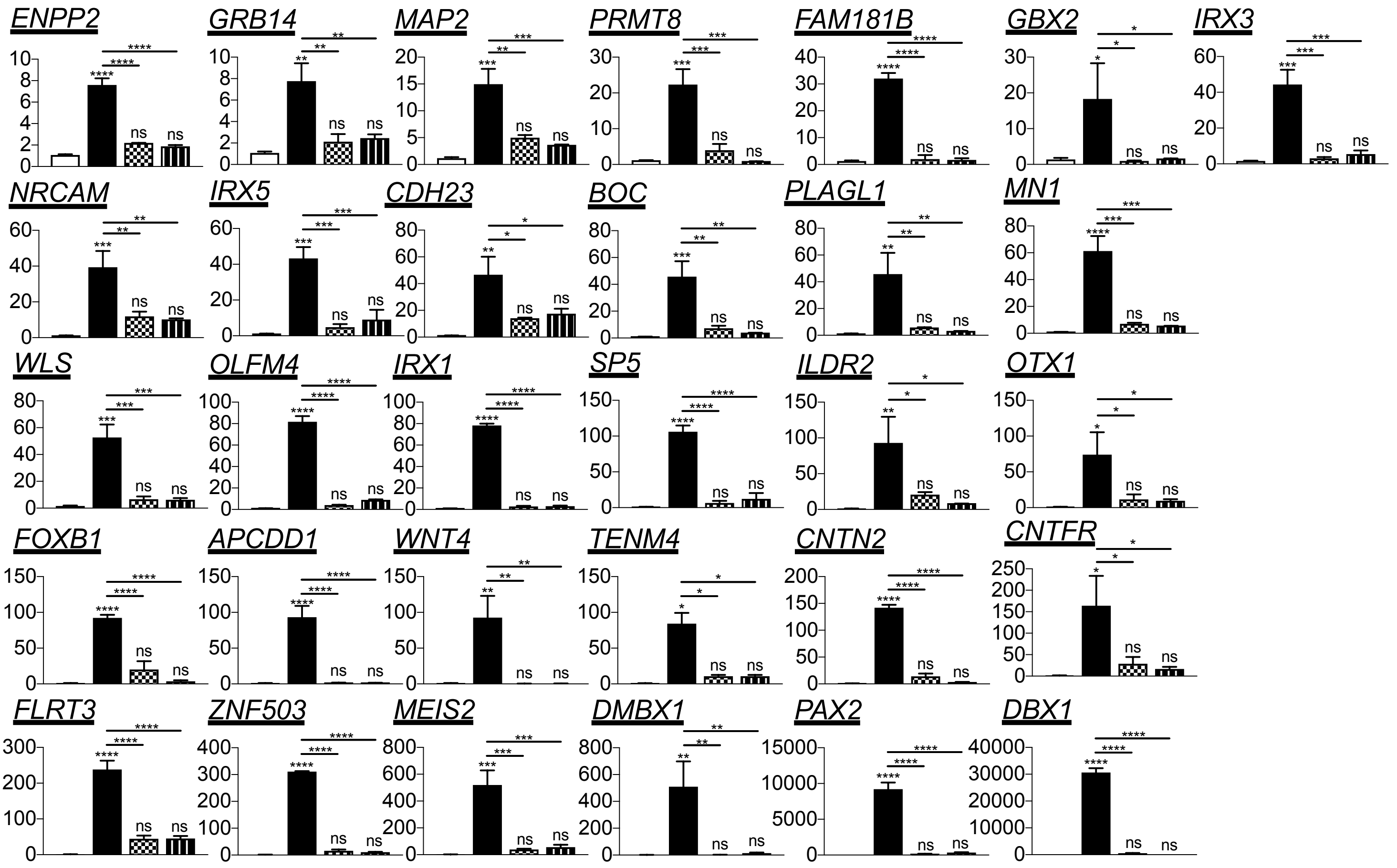


### F Neural crest enriched (N=13)

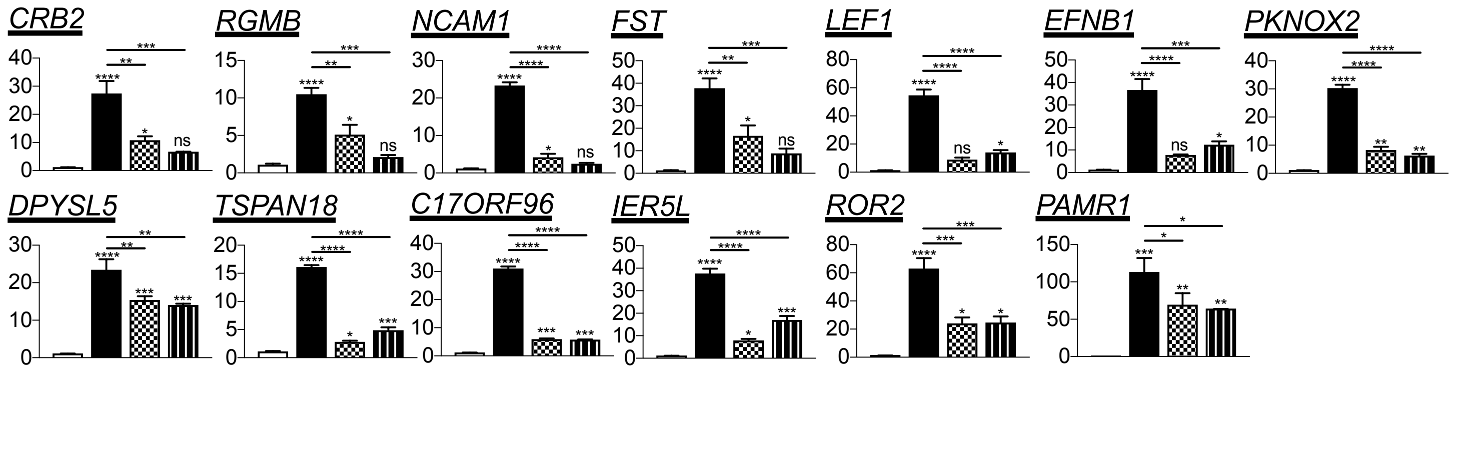

SUPPLEMENTAL FIG S1 RELATED TO DATA IN FIGURE 1

### G Neural crest biased (N=7)

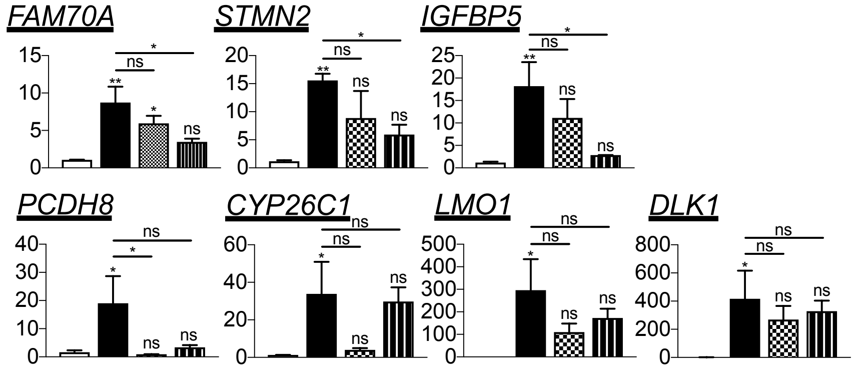


### H Broad ectoderm expression (N=7)

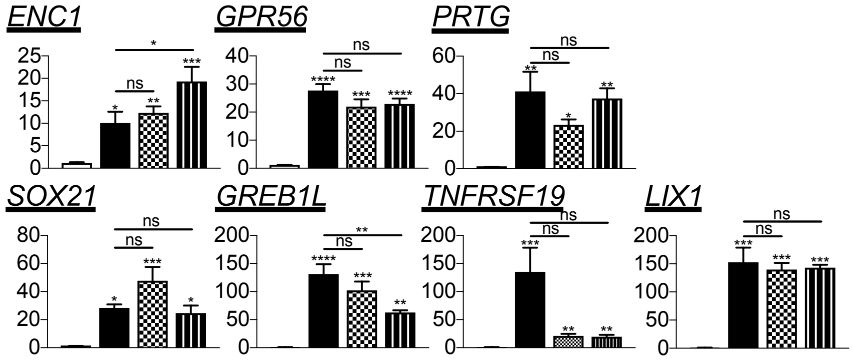


### I Highly variable expression during neural crest induction or not neural crest-specific (N=10)

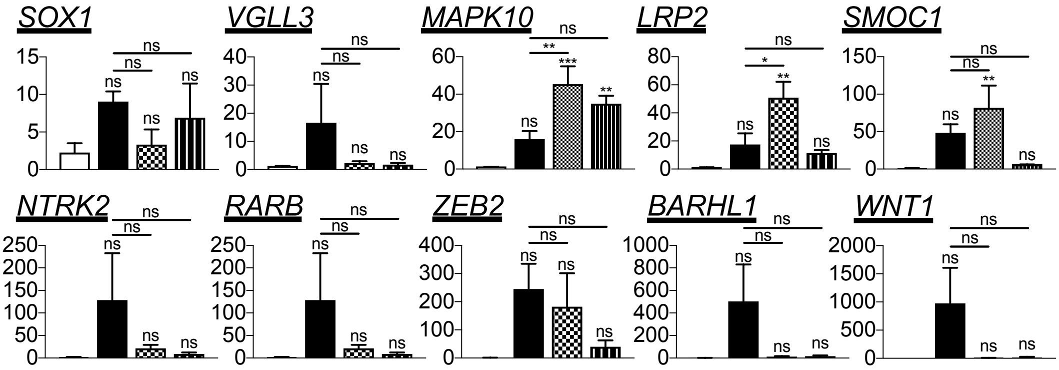

SUPPLEMENTAL FIG S2 RELATED TO DATA IN FIGURE 2

**A**

| | Upstream Regulator | Molecule Type | Predicted Activation State | Activation z-score | p-value of overlap |
|---|---|---|---|---|---|
| **Differentiation day 3** | **CTNNB1** | **transcription regulator** | **Activated** | **2.112** | **4.60E-27** |
| | beta-estradiol | chemical - endogenous | Inhibited | -3.122 | 3.26E-20 |
| | SOX2 | transcription regulator | | -1.921 | 5.19E-17 |
| | FGF2 | growth factor | | -1.146 | 9.82E-17 |
| | progesterone | chemical - endogenous | | -1.023 | 2.88E-15 |
| | WNT3A | cytokine | | -0.141 | 3.01E-15 |
| | GDF2 | growth factor | | -0.06 | 3.68E-14 |
| | TNF | cytokine | Inhibited | -2.527 | 1.82E-13 |
| | tretinoin | chemical - endogenous | | 0.144 | 1.66E-12 |
| | IFNG | cytokine | Inhibited | -3.779 | 1.85E-12 |
| | SMARCA4 | transcription regulator | | -1.783 | 8.15E-12 |
| | CREB1 | transcription regulator | Inhibited | -2.824 | 3.89E-11 |
| | POU5F1 | transcription regulator | | -0.135 | 6.04E-11 |
| | Vegf | group | | -0.001 | 6.15E-11 |
| | HGF | growth factor | | -1.301 | 1.28E-10 |
| | dexamethasone | chemical drug | Inhibited | -2.475 | 1.75E-10 |
| | TCF | group | | | 3.38E-10 |
| | Ngf | group | | -0.5 | 3.42E-10 |
| | Cg | complex | | -0.623 | 6.07E-10 |
| | Histone h3 | group | | | 6.42E-10 |
| **Differentiation day 5** | **CTNNB1** | **transcription regulator** | **Activated** | **3.415** | **1.36E-27** |
| | WNT3A | cytokine | | 0.676 | 6.45E-21 |
| | SOX2 | transcription regulator | | -1.531 | 1.95E-20 |
| | progesterone | chemical - endogenous | | -0.336 | 2.37E-20 |
| | tretinoin | chemical - endogenous | | 1.954 | 1.07E-18 |
| | TNF | cytokine | Inhibited | -2.458 | 2.02E-18 |
| | BMP4 | growth factor | Activated | 2.109 | 6.90E-16 |
| | beta-estradiol | chemical - endogenous | Inhibited | -2.244 | 1.34E-15 |
| | TGFB1 | growth factor | | 1.817 | 3.26E-15 |
| | estrogen receptor | group | | -1.981 | 1.24E-14 |
| | POU5F1 | transcription regulator | | -0.333 | 8.72E-14 |
| | BMP2 | growth factor | Activated | 2.512 | 3.75E-13 |
| | IFNG | cytokine | Inhibited | -3.818 | 4.69E-13 |
| | FGF2 | growth factor | | 0.804 | 6.52E-13 |
| | KLF4 | transcription regulator | | 1.148 | 6.88E-13 |
| | dexamethasone | chemical drug | | -0.994 | 8.73E-13 |
| | TWIST1 | transcription regulator | | 0.858 | 1.51E-12 |
| | decitabine | chemical drug | | -0.27 | 3.38E-11 |
| | SMARCA4 | transcription regulator | | -0.618 | 3.71E-11 |
| | STAT3 | transcription regulator | | -1.536 | 3.84E-11 |

**B**

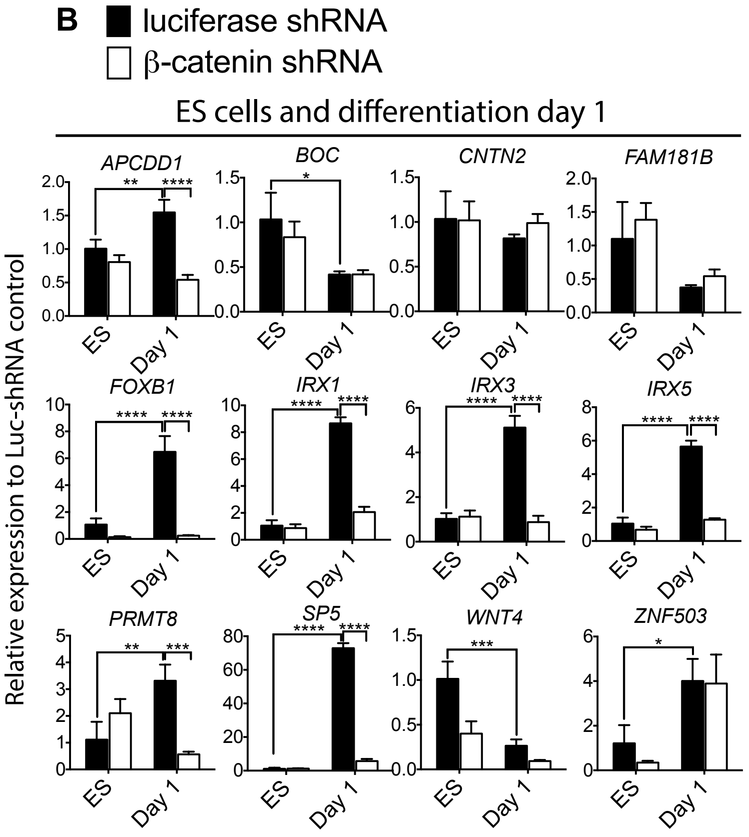


**C**

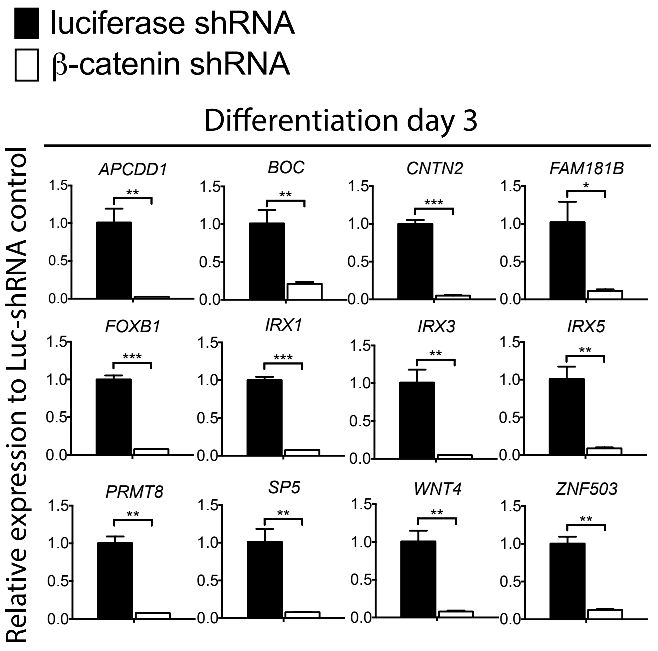

SUPPLEMENTAL FIG S3 RELATED TO DATA IN FIGURE 3

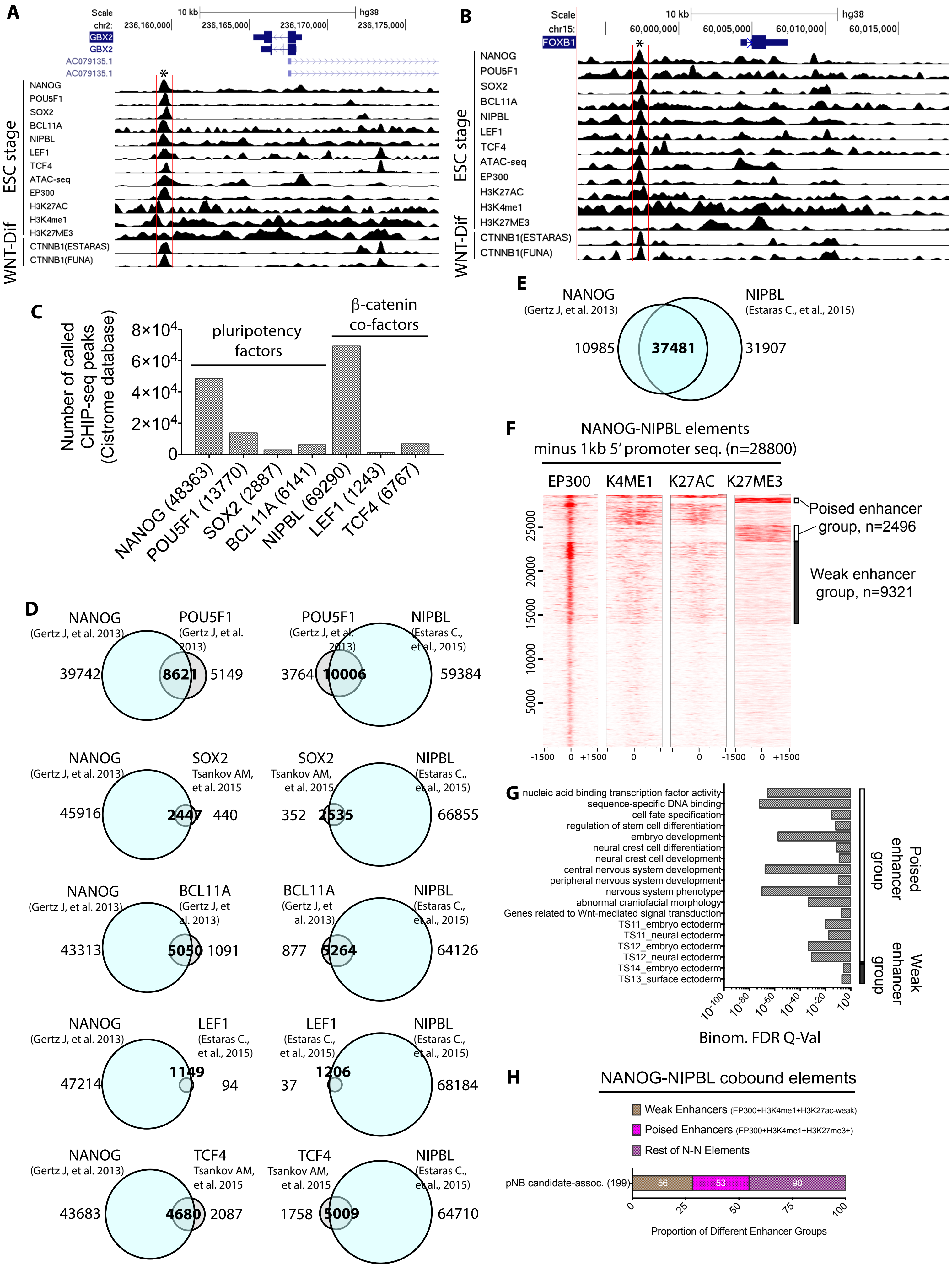

SUPPLEMENTAL FIG S4 RELATED TO DATA IN FIGURE 4

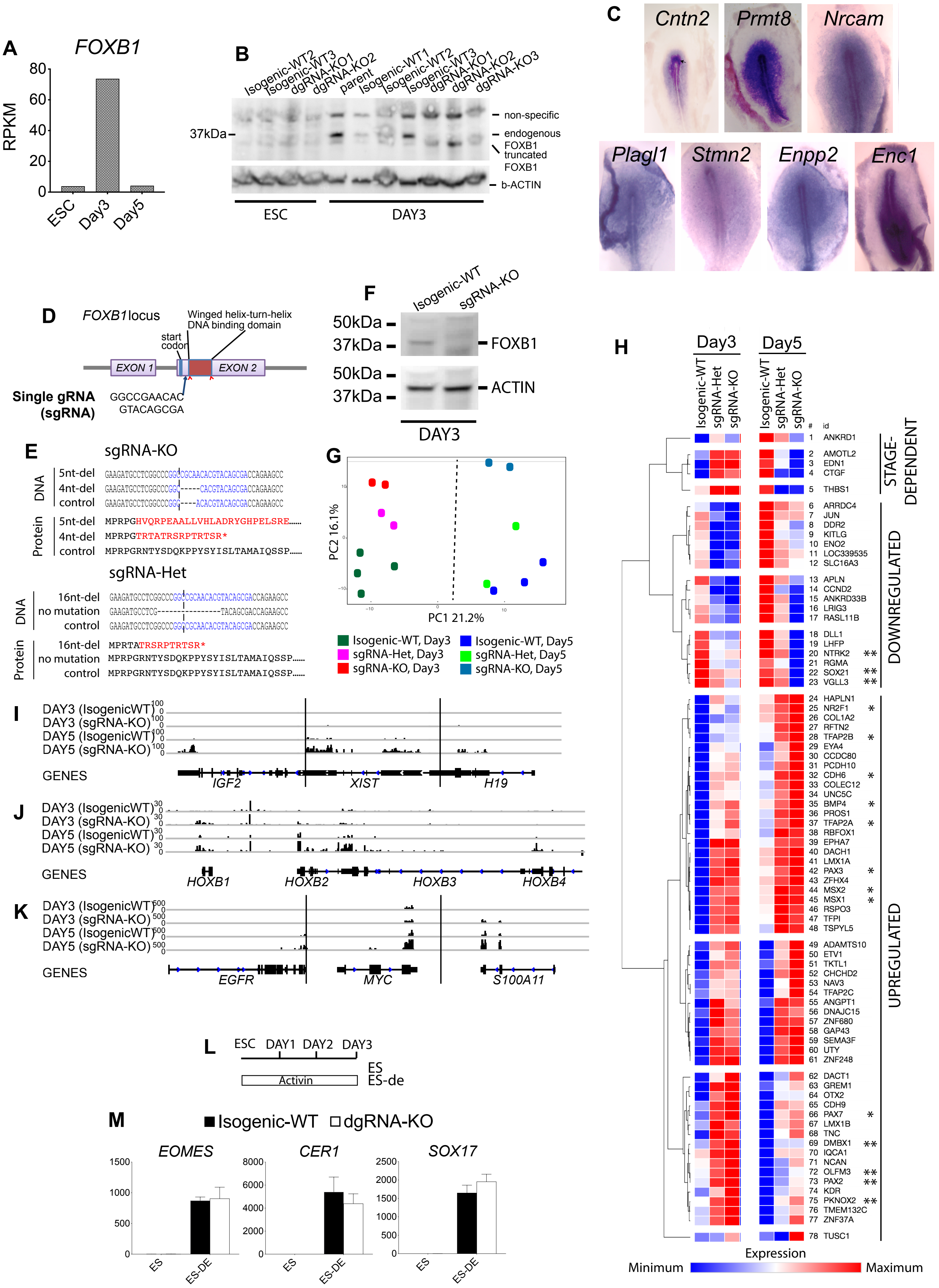

SUPPLEMENTAL FIG S5  RELATED TO DATA IN FIGURE 5

A

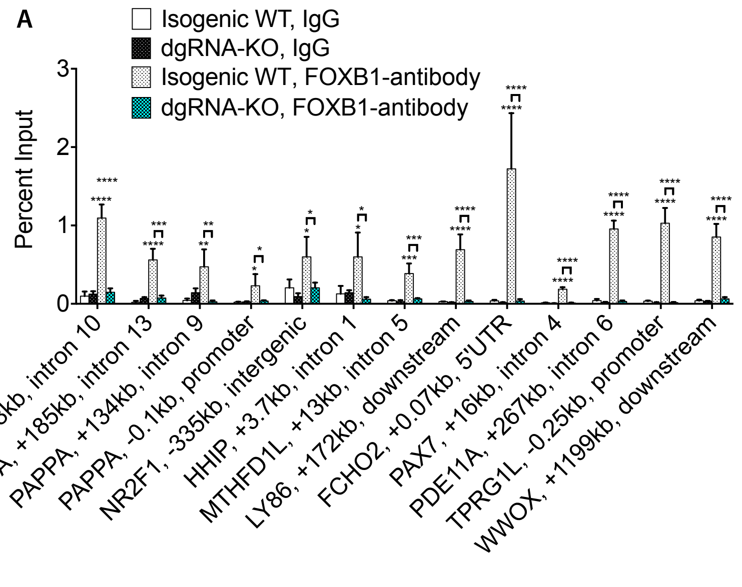

SUPPLEMENTAL FIG S6  RELATED TO DATA IN FIGURE 6

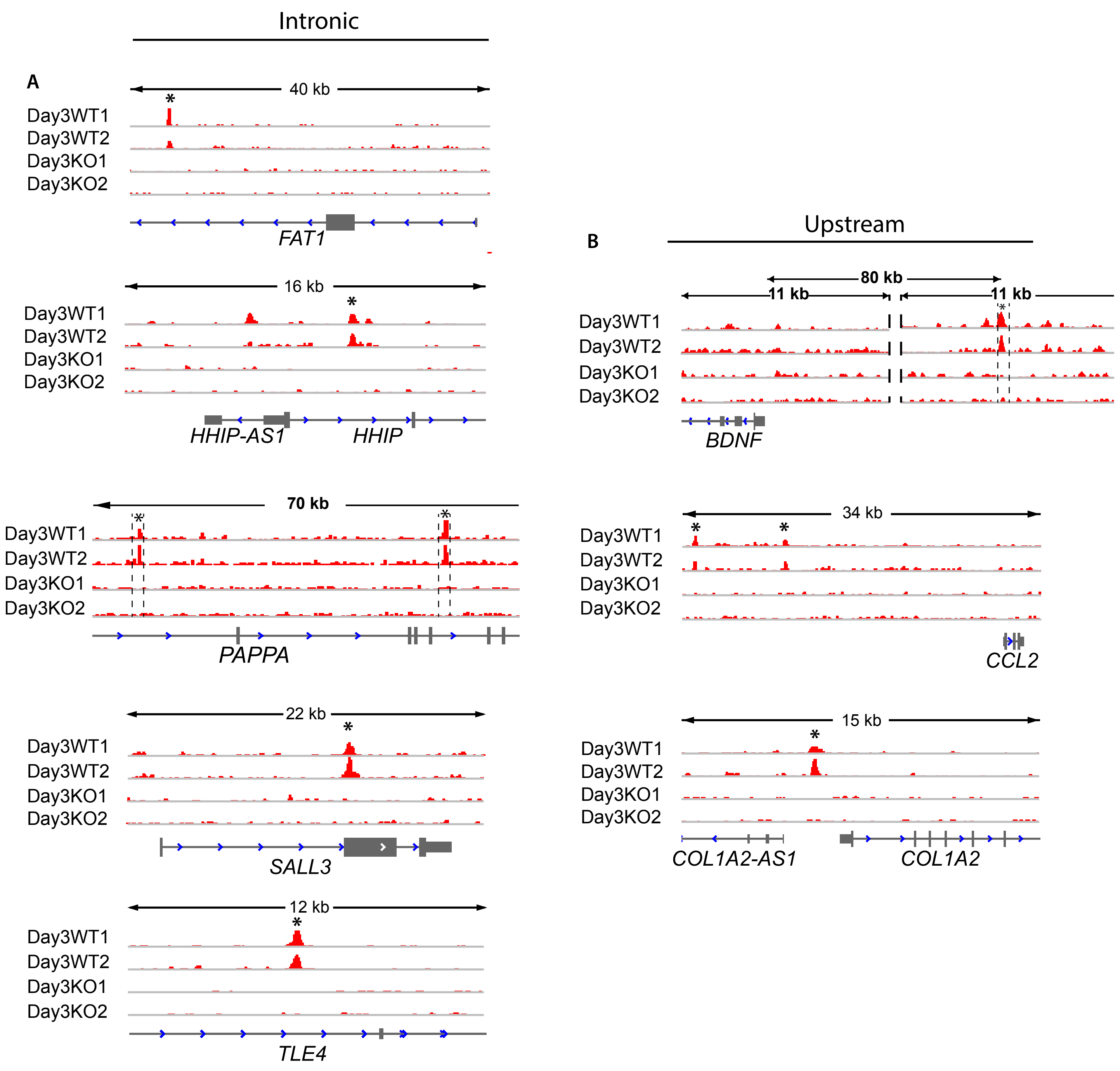

SUPPLEMENTAL FIG S7 RELATED TO DATA FROM FIGURE 7

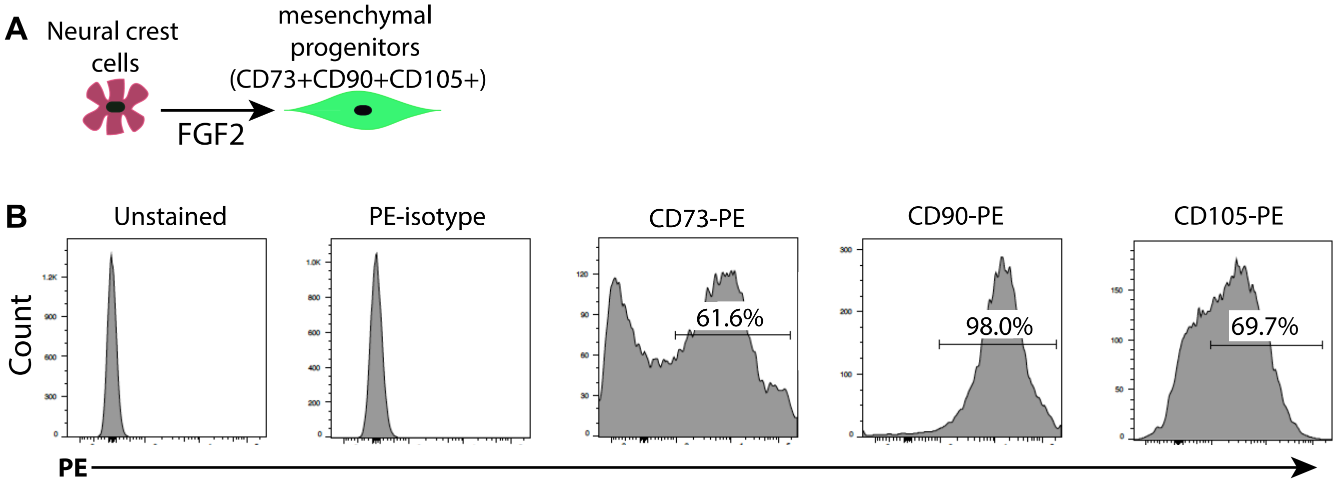
