## Supplemental text including methods and materials for "Pre-Border Gene Foxb1 Regulates the Differentiation Timing and Autonomic Neuronal Potential of Human Neural Crest Cells"

SUPPLEMENTAL MATERIALS

Methods and materials (page 2 to 10)

Supplemental figure and table legends (page 11 to 13)

Supplemental figures 1, 2, 3, 4, 5, 6 and 7 (pertaining to data presented in main figures 1, 2, 3, 4, 5, 6, and 7)

METHODS AND MATERIALS

Human ES and iPS Cell Growth and Maintenance

WA01 (H1) ES cells (WiCell), and Y6 iPS cells (Yale Stem Cell Center) from passages 35 to 60 were maintained as previously reported (Leung et al., 2016) on plastic surfaces coated with Matrigel (08-774-552, Fisher Scientific) in serum free medium (mTeSR1, STEMCELL^TM^ Technologies). Cultures were passaged every 6 to 7 days using Dispase (STEMCELL^TM^ Technologies) according to manufacturer’s instructions.

Neural Crest Cell Induction and Terminal Differentiation

Neural crest cell induction of human ES cells and iPSC was performed as previously described (Leung A. W., et al., 2016). For terminal differentiation, day 5 NC cells either in bulk or pre-sorted with anti-CXCR4 and anti-CD271 antibodies were replated onto matrigel-coated plates at 20,000 to 40,000 per cm^2^ in differentiation medium supplemented with 10 μM Y27632 for 24 hours. Differentiation medium for neuronal progenitors contained DMEM/F12 medium plus 2% B27 supplement, 1 x Glutamax, 3 μM CHIR 99021 (Tocris), 1 μM SU4301, and 2.5 μM DAPT. Differentiation medium for mesenchymal progenitor contained DMEM/F12 medium plus 2% B27 supplement, 1 x Glutamax, and 20 ng/mL FGF2 (R&D systems). Differentiation was carried for 6 days with medium change at every other day.

Neurectoderm, Mesoderm and Endoderm Induction

For neurectoderm induction, human ES cells were plated at 20,000 cells per cm^2^ on matrigel-coated plates in differentiation medium containing DMEM/F12 medium plus 2% B27 supplement, 1 x Glutamax, 100 nM LDN-193189 (Tocris) and 10 μM SB431542 (Tocris). For mesoderm and endoderm induction, human ES cells were plated at same density on matrigel-coated plates in differentiation medium containing RPMI medium plus 2% B27 supplement, 1 x Glutamax, plus 20 ng/mL FGF2 (R&D system)(mesoderm) or 100 ng/mL Activin (ConnStem)(endoderm) respectively.

Transcript Analyses

Cells were dissolved in TRIzol® reagent (Life Technologies) and total RNAs were extracted according to manufacturer’s instructions. Total RNA (0.5 to 1 μg) was reverse-transcribed using SMART MMLV reverse transcriptase (Clontech Laboratories, Inc.). Real time polymerase chain reaction (qPCR) was set up using the iTaqTM universal SYBR® Green supermix (BIO-RAD) with primer concentration at 160 nM and reactions were run on either a CFX96 or a CFX384 Touch™ Real-Time PCR Detection System (BIORAD). Forty cycles of reactions were performed followed by running melting curves to confirm specificity of each primer sets. Six or more biological replicates from at least two independent experiments were measured unless otherwise specified. Primer sequences are listed in table S6.

Lentivirus Production and Generation of Stable Human Pluripotent Stem Cell Lines

Lentivirus generation and infection for WA01 human ES cells to produce luciferase-shRNA and β-catenin shRNA were performed as described previously (Leung et al., 2016).

Generation of CRISPR-Cas9 Edited Human Pluripotent Stem Cells Lines

sgRNA-Het, dgRNA-KOs, and FOXB1-E1/E2-KOs were generated using an improved transfection and selection protocol using *Cas9* mRNA (A.W.L. and Dr. Yan-Ru Lou, University of Helsinki, Finland, manuscript in preparation). sgRNA-KO was generated by lipofection of Cas9-protein and FOXB1-crRNA:tracrRNA duplex. In brief, 1 μg of *in vitro* transcribed *Cas9* mRNA from pCS2-Cas9 (Dr. Takamoto) and 10 pmol each of crRNA:tracrRNA complex (IDT) were transfected into human ES cells. crRNA sequences for targeting *FOXB1* enhancer E1, *FOXB1* translational start site (sgRNA) and *FOXB1* DNA binding domain sequence (dgRNA) were TGCTAGAGCGCCCAAAGCAG / CATACTACACACGTTCACCC, TCGCTGTACGTGTTGCGGCC and GTACATCTCGCTGACCGCTA / GGTGCTTAAGTCCGACCACC respectively. Single clones were expanded and genotyped using primer sets forward: CGAGAGAAGGAGGGTAGAGG / reverse: GGTAAATCCCGGACAGTAGG, forward:CGCAACTTGAAGCAACTTTA / reverse: TTGATGAAGCAGTCGTTGAA and forward: AGAAGAGGGCGAGGAAGAAG / reverse: GCGATGATGTTCTCGATGG for FOXB1-E1 enhancer, FOXB1 sgRNA and FOXB1 dgRNA respectively. Mutations were verified by PCR genotyping, Sanger sequencing and/or western blotting. Additional verification was performed on mutants generated using single crRNA by TA-cloning using TA Cloning^TM^ kit (Thermo Fisher Scientific).

siRNA mediated Knockdown

siRNA mediated knockdown was performed using SMART Pool siRNA for FOXB1 (Dharmacon cat.# L-008906-00) and non-targeting control siRNA (Dharmacon cat.# D-001810-10) transfected with Lipofectamine RNAiMAX (Thermo Scientific cat. # 13778075). Reverse transfection was performed as per manufacturers instructions. Briefly, siRNA/RNAiMAX mixture was pre-plated into the 96 well plate in technical replicate for each biological replicate. Human ES cells were plated in NC induction media as described above into each of the wells containing siRNA/RNAiMAX mixture. For each well 1.2 picomole of siRNA was used with 0.1 μl of Lipofectamine RNAiMAX reagent in total 20 μl of OptiMEM medium. Medium was changed every 24 hours and cells were harvested at day 5 for RNA extraction and immunofluorescence.

Western Blotting

Approximately 1 million cells were collected by Accutase digestion. The cells were spun down at 500g for 5 min and dissolved in 100 μL Pierce® RIPA buffer (Thermo Scientific) plus 33 μL 3:1 Laemmlli buffer:β-mecaptoethanol. Cell solution was then sonicated at a 30s-on/30s-off scheme at high intensity for 10 min using Bioruptor^TM^ UCD-700. The sonicated solution was spun down at 14,000 rpm for 5 min at 4^o^C to remove cell debris. The samples were then heated at 95^o^C for 10 min. Samples can be stored at -20oC. Samples were run on mini-protean TGX^TM^ Precast Gels (4-20% gradient gel, Biorad) at 120V for 45 to 60 min. Proteins were then transferred to an activated PVDF membrane (1620177, Biorad) in a Biorad electrophoresis transfer system at 4^o^C for 70 min at 240 mA. The membrane was then blocked in 5% milk in PBS. 1:2000 goat anti-human FOXB1 antibody (abcam) in 1% BSA in PBS was used to incubate membrane overnight at 4^o^C. After overnight incubation, membrane was washed at least 3 times in 0.05% Tween-20.  The membrane was then probed with an anti-goat-HRP (1:3000). Signals were then developed using ECL kit and images taken on Syngene Bio Imaging machine using Genesnap software. After detection of FOXB1 signal, membrane was stripped in Restore^TM^ Plus Western Blot Stripping Buffer (Thermo Scientific) for 8 min at RT and washed PBST. 1:3000 anti-ACTIN-HRP for 40-60 min in 5% milk was then added and membrane washed with PBST. Signals were then developed as described above.

Chromatin Immunoprecipitation

Approximately 10 million cells were dissociated into single cells using Accutase. Cells were washed once and fixed in 5 mL DMEM/F12 medium with 1% paraformaldehyde (electrical) for 7.5 min at RT with rotation. Fixation was stopped with addition of 625 μL 1M glycine followed by 5 min rotation at RT. Cells were then spun down at 4^o^C at 500g for 5 min, followed by two washes of ice-cold PBS with protease inhibitor. Cell pellets were snap frozen at -80^o^C until use. At day of ChIP, cell pellets were resuspended in 1 mL Lysis Buffer 1 containing 50 mM Hepes-sKOH, pH7.5, 140 mM NaCl, 1 mM EDTA, 10% Glycerol, 0.5% NP-40, and 0.25% Triton X-100 followed by mixing at 4^o^C for 10 min. Cell pellets were spun down at 1400g for 5 min and resuspended in 1 mL Lysis Buffer 2 buffer containing 10 mM Tris-HCl, pH8.0, 200 mM NaCl, 1 mM EDTA, and 0.5 mM EGTA. Following another 10 min incubation at 4^o^C, cell pellets were again spun down at 1400g for 5 min and resuspended in 200 to 300 μL Lysis Buffer 3 containing 10 mM Tris-HCl, pH8.0, 100 mM NaCl, 1 mM EDTA, 0.5 mM EGTA, 0.5% Na-Deoxycholate and 0.5% N-lauroylsarcosine. Cells were then sonicated using Covaris S200 machine with the following parameters: peak power=120, duty factor=2.0, cycle/burst=200, temperature=5^o^C and scale=10 min and a power output at 2.1. The sonicated cell solution was topped up to 600 μL with Lysis buffer 3 plus 60 μL 10% Triton X-100, incubated for 15 min with rotation at 4^o^C and spun down at 20,000g for 15 min. The supernatant was then divided into input tube (60 μL), IgG antibody control tube (300 μL) and target antibody tube (300 μL). 5 μg of anti-NANOG (AF1997, R&D systems) and anti-FOXB1 antibodies (ab5274, abcam), and 1.5 μg of anti-H3K27ac antibody (rabbit antibody) were added as target antibodies with equivalent amount of species-specific antibodies for IgG antibody controls. After overnight incubation at 4^o^C, antibodies were pulled down using Protein G dynabeads (10 μL per 1 μg antibody) and dynabeads were extensively washed with RIPA buffer containing 50 mM Hepes-KOH, pH7.6, 500 mM LiCl, 1 mM EDTA, 1% NP40 and 0.7% Na-Deoxycholate with rotation at 4^o^C for 3 to 4 hours. After last RIPA wash, rinsed and washed once with TE+NaCl buffer containing 50 mM Tris-HCl, pH8.0, 10 mM EDTA and 50 mM NaCl. DNA was then eluted in 125 μL elution buffer containing 50 mM Tris-HCl, pH8.0, 10 mM EDTA and 1% SDS at 65^o^C for 20 min. Eluted DNA was then transferred to a 1.5 mL DNA LoBind tube (eppendorff) for overnight incubation at 65^o^C for reverse cross-link. Input tube was also reverse cross-linked in 2 to 3 volumes of elution buffer. 1 μL of RNase A1/T mix (Thermo Scientific) was added to reverse cross-linked DNA the next day to digest endogenous RNAs and the solution was incubated at 37^o^C for 1 hr. This is followed by addition of 3 μL of proteinase K and incubation for 1 to 2 hr at 56^o^C. The solution was then phenol-chloroform extracted once followed by DNA extraction using mini-Elute kit (28004, Qiagen). DNA was eluted in 50 to 60 μL with EB buffer for downstream qPCR analysis or ChIP-seq library construction.

Sequencing Library Preparations

*RNA-seq libraries*

Libraries were prepared from poly-adenylated mRNAs using a custom kit based on Illumina’s mRNA-seq kit (Illumina, part # RS-122-2101) and sequenced by Illumina HiSeq2000 or Hiseq2500 machine for 76 base pair read lengths in the Yale Center for Genome Analysis (YCGA) or Yale Stem Cell Center Genomics and Bioinformatics Core.

*ChIP-seq libraries*

Approximately 10 – 15 ng ChIP DNA (around 15 μL) for each sample was used for ChIP-seq library construction following manufacturer instructions for NEBNext® UltraTM II DNA Library Prep Kit for Illumina (New England Biolabs). In brief, ChIP DNA was ended repaired at 20^o^C for 30 min followed by incubation at 65^o^C for 30 min. Ligation of adaptor (NEBNext Adaptor for Illumina) was then performed at 20^o^C for 15 min followed by U excision at 37^o^C for 15 min. Adaptor ligated and U-excised DNA was then cleaned up using AMPure XP beads (Beckman Coulter), washed twice with freshly prepared 80% ethanol and eluted in 17 μL of 10 mM Tris-HCl. PCR amplification was used to add the following indices GTCCGC, GTGAAA, GTGGCC, GTTTCG, CGTACG, and GAGTGG (NEBNExt Index 18 to 23) to the cleaned up DNA followed by another clean up step with AMPure XP Beads as described above. DNA was eluted in 30 μL 0.1X TE. The amount and quality of DNA was measured on Qubit 3.0 Fluorometer (Life Technologies) using Qubit™ dsDNA HS Assay Kit and Bioanalyzer (Agilent) using High Sensitivity DNA chip.

Sequencing Read Processing, Mapping and Analyses

*RNA-seq*

Three different RNA-seq experiments are conducted in this study. Sequences and expression profiles for each sample, together with the data processing details can be found in NCBI’s Gene Expression Omnibus (GEO) under accession number GSE125145. Briefly, sequencing reads were trimmed for low quality bases. Trimmed reads were mapped to the human reference genome (hg19) using Tophat v2.1.1 or STAR. Alignments were then processed by Cuffdiff (Cufflinks v2.2.1) to obtain differential gene expression providing gene model annotation and the genome sequence file for detection and correction of sequence-specific bias arise from the use of random hexamer during library preparation. For downstream analysis, tab-delimited text files include RPKMs, fold changes, and p-values for each condition in Cuffdiff output format were used.

*ChIP-seq*

For each sequenced read, we trimmed the first 6 nucleotides and the last nucleotides at the point where the Phred score of an examined base fell below 20 using in-house scripts. If, after trimming, the read was shorter than 45 bp, the entire pair was discarded. Trimmed reads were mapped to the human reference genome (hg19) using BWA-MEM v0.7.17 using default parameters. Only reads with mapping quality scores equal or higher than 20 were kept. Duplicated read pairs were removed to allow only unique fragments using Picard MarkDuplicates v2.8.2 (http://broadinstitute.github.io/picard/). Peak and motif finding was performed using HOMER v4.10 with default parameters for transcription factors. bigwig for each sample (bw) and tabulated text file with Homer peakcaller output (peak) were used for analysis. Motif from called peaks were also called using Seqpos software (Version: 0.590) on Cistrome/galaxy with *de novo* motif and known motif analyses.

Visualization of Expression Data with Heat Maps

Visualization of RNA-seq expression data on heat maps was performed using Morpheus software (https://software.broadinstitute.org/morpheus/).

Public Dataset Acquisition

Bed and bigwig files for NANOG (GSM803437), NIPBL (GSM1579363), SOX2 (GSM1505768), POU5F1 (GSM803438), BCL11A (GSM803396), LEF1 (GSM1579343), TCF4 (GSM1505791), CTNNB1 (GSM1412243; GSM1579346), H3K4me1 (GSM733782), H3K27ac (GSM733718), H3K27me3 (GSM733748), and EP300 (GSM602291) were downloaded from the Cistrome project website (http://cistrome.org/) with build hg38. Conversion of builds for bed files was performed using *Liftover* from UCSC genome browser.

Embryo Preparation

Fertilized *Gallus gallus* eggs were obtained from Hardy’s Hatchery (Massachusetts, USA) and incubated at 37°C until it was estimated that they had reached the required Hamburger-Hamilton stage of development. Once at the appropriate stages for *in situ* hybridization, the embryos were harvested and dehydrated in 100% methanol. Embryos were washed in Ringer’s solution and fixed in 4% Paraformaldehyde at 4°C overnight.

Whole Mount *In Situ* Hybridization and Probe Generation

Whole mount *in situ* hybridization was carried out as previously described (Basch et al., 2006). The *Foxb1* antisense probe was generated by amplifying chick cDNA with Foxb1 forward primer ACTTTAAGATCTCCGCGGGT and reverse primer aagcttTAATACGACTCACTATAGGGAGAATGTTCTCGATGGCGAAGGG in a PCR reaction. NEB Taq 2X Master Mix (M0270) was utilized for the PCR reaction and the product size was 547 bp. The PCR product was excised from the gel and purified using the Zymo Gel DNA Recovery Kit. Following this, *in vitro* transcription was performed using a T7 polymerase to label RNA with digoxigenin-UTP (Roche DIG RNA Labeling Mix, 10X Conc.) The antisense probe was then purified using an Illustra Microspin G-50 Column. Whole mount in situ hybridization was performed on chick embryos that were collected in Ringer’s Solution and fixed in 4% Paraformaldehyde overnight. The embryos were dehydrated through a series of methanol solutions and stored at -20°C. The embryos were hybridized with the Foxb1 antisense probe at 65°C overnight. The probe was then removed and the embryos were exposed to several post-hybridization washes. The embryos were left overnight in alkaline phosphatase-anti-DIG antibody at 4°C. The following day NBT/BCIP was utilized to develop the ISH signal. Probes for other pB candidates were prepared similarly as described above.

Embryo Sectioning

The embryos were incubated for several hours in 5% sucrose, 15% sucrose, and 7.5% gelatin solutions before sectioning. The embryo was then embedding in gelatin and cryo-sectioned at 10 µm for stages HH4 and HH6 and 15 µm for HH8.

Pathway Analysis

Ingenuity pathway analysis software, licensed to Yale University (Version 01-07), was used to perform upstream regulator, canonical pathway and interacting partner analyses. Gene ontogeny analysis was also performed using GREAT software (http://great.stanford.edu/public/html/).

Statistical Analyses

Chart drawing and statistical analyses were performed using the GraphPad software Prism (version 7.0f). Asterisks were used to indicate statistical significance with * p-values ≤ 0.05, ** p ≤ 0.01, *** p ≤ 0.001, **** p ≤ 0.0001 unless otherwise indicated. ‘ns’ indicates no significant difference detected. Unpaired T-tests or one-way ANOVA tests with Tukey or Bonferroni corrections for multiple comparisons were respectively performed for comparing 2 or more means unless otherwise specified in figure legends. Association analyses were performed using the IntersectRegions function in the open software Useq package (http://useq.sourceforge.net).

SUPPLEMENTAL FIGURE AND TABLE LEGENDS

**Supplemental Figure 1.** Data related to Figure 1. (B) Diagram displaying the number of differentially expressed genes (p<0.05) for each of the indicated comparisons. (B-F) K-means clustering of the RPKM values of genes grouped according to their known biological functions or gene clusters in the ESC, day 3 and day 5 transcriptomes. Relative intensities were displayed with blue being lowest in expression level to red being highest. (D) Differentiation schemes for ES, NC, NE and NNE progenitor cell types. (E-I) qPCR quantification of selected pB candidates categorized according to their expression levels relative to ES, NE, and NNE. Multiple comparisons were carried out by Fisher’s LSD test. Asterisks directly above bars indicate comparison between control ES group either with NC, NE or NNE group.

**Supplemental Figure 2.** Data related to Figure 2. (A) Upstream regulator analysis on genes that were differentially expressed in day 3 and day 5 transcriptomes using IPA software. (B-C) qPCR data comparing control (luciferase shRNA) against knockdown (*β-catenin* shRNA) in ES and day 1 (B), and day 3 (C) cultures.

**Supplemental Figure 3.** Data related to Figure 3. *GBX2* (A) and *FOXB1* (B) loci with embedded published ChIP-seq and ATAC-seq data sets in bigwig formats were displayed on UCSC browsers. ‘ESC stage’ indicates the sequencing data were derived from human ES cells. ‘WNT-Dif’ indicates the data were from differentiating cells treated with CHIR 99021. (C) Bar chart displaying the number of peaks called from each of the indicated published ChIP-seq data sets (number in brackets) grouped into pluripotency factors (NANOG, POU5F1, SOX2, BCL11A) and β-catenin co-factors (NIPBL, LEF1, TCF4). (D-E) Venn diagrams showing the relative proportion of overlaps and the number of overlapping peaks for the indicated comparing peak sets. (F) K-means clustering of EP300, H3K4me1, H3K27ac and H3K27me3 signals on NANOG-NIPBL co-bound elements. Poised enhancers were defined as EP300+H3K4me1+H3K27ac-H3K27me3^strong^ while weak enhancers as EP300+H3K4me1+H3K27ac^weak^H3K27me3^weak^. (G) Bar chart showing gene ontogeny terms that were significantly associated with NANOG-NIPBL co-bound elements present in the poised and weak enhancer categories. (H) Proportion and number of NANOG-NIPBL elements that were associated with the 68 pB candidates.

**Supplemental Figure 4.** Data related to Figure 4. (A) Plot for RPKM values for human FOXB1 gene from time course RNA-seq described in Fig 1 and Fig 4A. (B) Western blot on ES cell and differentiation day 3 cultures of parent, isogenic control and dgRNA-KO cell lines using anti-FOXB1 antibody. (C) Images of Chick HH4 embryos performed with in situ hybridization using the indicated gene probes. Arrow within *Cntn2* in situ image panel indicates the node structure. (D) CRISPR targeting schematic for FOXB1 locus using a single gRNA. (E) Sanger-sequencing data from each allele of the two *FOXB1* mutant cell lines (*DNA*). Predicated amino acid sequences are displayed as well (*protein*). (F) Western blot of day 3 cultures derived from an isogenic control and the sgRNA-KO cell line using anti-FOXB1 and anti-ACTB antibodies. (G) Principal component analysis of RNA-seq data from isogenic control, sgRNA-Het and sgRNA-KO cell lines collected at day 3 and day 5. (H) K-means clustering of the RPKM values of selected genes clustered according to expression profiles in isogenic control and *FOXB1* mutant cell lines. (I-K) Bigwig files of day 3 and day 5 isogenic wildtype and FOXB1 sgRNA-KO cell line displaying transcript signals for selected gene groups on IGV software. (L) Differentiation and sample collection schematic for endoderm competency test. (M) *EOMES*, *CER1,* and *SOX17* RT- qPCR of cells differentiated according to schematic in panel L.

**Supplemental Figure 5.** Data related to Figure 5. (A) ChIP-qPCR data on FOXB1-bound elements identifiesd from FOXB1 ChIP-seq. Asterisks directly above ‘Isogenic-WT, FOXB1 antibody’ bar indicates significant differences detected between IgG and FOXB1 antibody groups. Distance from transcriptional start sites of closest genes was also shown for each of the elements. One-way ANOVA with Sidak correction for multiple testing was performed on individual FOXB1-bound regions.

**Supplemental Figure 6.** Data related to Figure 6. IGV tracks displaying ChIP-seq signals of 2 wildtype ES cell samples and 2 dgRNA-KO samples for loci containing gene transcripts that are upregulated during human neural crest induction. Selected FOXB1-bound regions found within intronic sequences (A) or upstream of transcription start sites (B) are displayed. Asterisks indicate location of called peaks.

**Supplemental Figure 7.** Data related to Figure 7. (A) Differentiation schematic for mesenchymal progenitors from NC cells. (B) Flow cytometry analysis of controls and antibody-stained cells using anti-CD73-PE, anti-CD90-PE and anti-CD105-PE antibodies.

**Table S1.** RPKM values, fold changes, and p-values from all mapped transcripts in ESC, day 3 and day 5 RNA-seq data set described in Figure 1.

**Table S2.** RPKM values, fold changes, and p-values from all mapped transcripts in Luc-shRNA and CTNNB1 shRNA RNA-seq data set described in Figure 2.

**Table S3.** Statistics of intersecting peaks set analysis for pluripotency factors and b-catenin co-factors.

**Table S4.** RPKM values, fold changes, and p-values from isogenic wildtype, FOXB1 sgRNA-Het and FOXB1 sgRNA-KO cell line transcriptomes described in Figure S4.

**Table S5.** FOXB1 direct targets deduced from FOXB1-CHIP-seq data within a 50kb window from target gene transcription start sites.

**Table S6.** Primer information for transcript and ChIP-qPCR analyses.
